# Molecular and cellular insight into *Escherichia coli* SslE and its role during biofilm maturation

**DOI:** 10.1101/2021.02.07.430137

**Authors:** Paula M. Corsini, Sunjun Wang, Saima Rehman, Katherine Fenn, Amin Sagar, Slobodan Sirovica, Leanne Cleaver, Charlotte J. C. Edwards-Gayle, Giulia Mastroianni, Ben Dorgan, Lee M. Sewell, Steven Lynham, Dinu Iuga, W. Trent Franks, James Jarvis, Guy H. Carpenter, Michael. A. Curtis, Pau Bernadó, Vidya C. Darbari, James A. Garnett

## Abstract

*Escherichia coli* is a Gram-negative bacterium that colonizes the human intestine and virulent strains can cause severe diarrhoeal and extraintestinal diseases. The protein SslE is secreted by a range of pathogenic and some commensal *E. coli* strains. It can degrade mucins in the intestine, promotes biofilm maturation and it is a major determinant of infection in virulent strains, although how it carries out these functions is not well understood. Here we examine SslE from the *E. coli* Waksman and H10407 strains and reveal that it has a novel and dynamic structure. In response to acidification within mature biofilms we show how SslE forms a unique functional aggregate that interacts with cellulose and regulates the distribution of exopolysaccharides in macrocolony biofilms. Our data indicates that the spatial organization of SslE polymers and local pH are critical for biofilm maturation and SslE is a key factor that drives persistence of SslE-secreting bacteria during acidic stress.

## Introduction

*Escherichia coli* is a primary colonizer of the lower intestinal tract of humans and other warm‐blooded animals. While many strains are considered beneficial to the host and help to maintain a healthy immune system, virulent strains are the cause of severe diarrhoeal diseases, including haemorrhagic colitis, and extraintestinal diseases such as neonatal meningitis, urinary tract infections, sepsis and pneumonia^1^. A wide range of pathogenic *E. coli* strains, and some commensals, use a *Vibrio*-like type II secretion system (T2SS) to translocate the protein SslE across their outer membrane and onto their extracellular surface^2, 3, 4, 5, 6, 7^. These include the Waksman (W), enterotoxigenic (ETEC), enterohemorrhagic (EHEC), enteropathogenic (EPEC), enteroaggregative (EAEC), enteroinvasive (EIEC) and neonatal meningitis *E. coli* (NMEC) strains. SslE is required for full virulence in a rabbit model of EPEC infection^4^ and as a surface exposed antigen, SslE has shown great promise as a broadly protective vaccine candidate against a wide range of *E. coli* pathotypes^3, 6, 8, 9^.

SslE interacts with mucosal membranes in the host intestine where it can degrade mucins^6, 10, 11, 12^, a family of heavily *O*-linked glycosylated proteins and the primary constituents of mucus^13^. This provides nutrients during bacterial growth but also enables these *E. coli* strains to penetrate the gut mucosa to access host cells for efficient colonisation and targeting of toxins/effectors. Furthermore, SslE is also important for mediating maturation of EPEC biofilms^4^; microbial aggregations encased within a self-produced extracellular matrix comprised of exopolysaccharides, adhesive proteins and nucleic acids^14^. When *E. coli* is released into the environment through faeces or wastewater effluent it can survive for long periods within complex biofilm communities^15, 16^, and these are fundamental for both the environmental ecology of *E. coli* but also for successful colonisation of the intestinal tract^17^. However, the specific molecular mechanisms that SslE uses to promote ecology and disease are not well understood.

SslE is a ~165 kDa lipoprotein composed of an N-terminal periplasmic localization sequence and lipobox motif, an unstructured ~5 kDa region, a ~110 kDa region with no significant primary sequence homology to any other known protein, and a ~50 kDa M60-like aminopeptidase domain at its C-terminus^18^ (**Fig. 1a**). M60-like domains are metalloproteases that contain a zinc binding HExxH motif and an additional conserved catalytic glutamate residue, which cleaves the peptide backbone of mucin-like substrates. These and other related enzymes have been identified in both prokaryotic and eukaryotic microbes that interact with host mucosal membranes^18^ and structures of proteoglycan complexes suggest that interactions with both the mucin peptide and *O*-linked carbohydrate sidechains are important for specificity^19, 20^. The peptidase activity of the SslE M60 domain is effective for the degradation of major mucins of the intestine (e.g. MUC2, MUC3, MUC5AC)^6, 10, 11, 12^, however, very little is known as to how SslE interacts with these mucins or the function of the remaining SslE sequence.

**Fig. 1.**
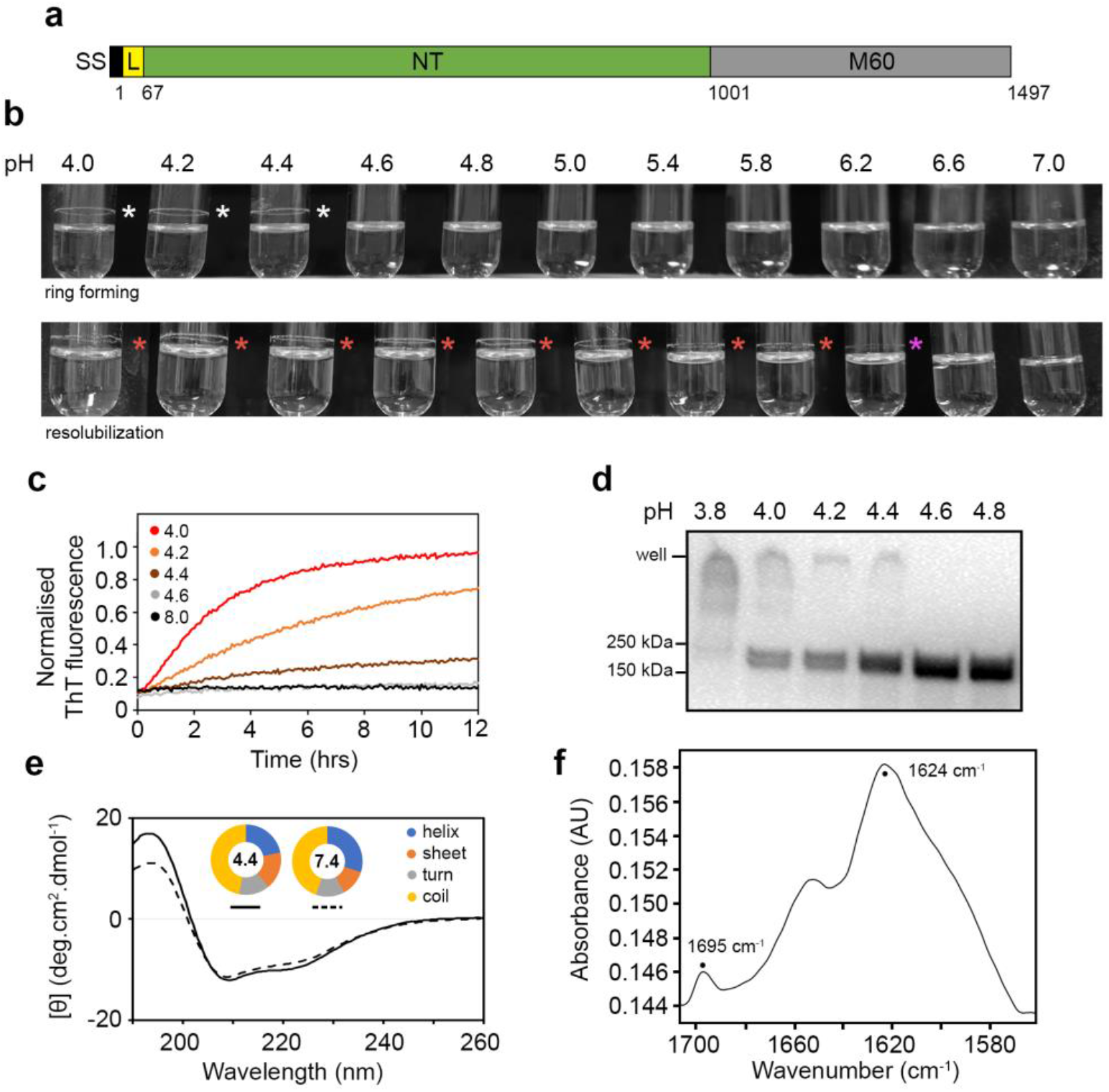
Characterization of SslE aggregates. **a,** Schematic of SslE from *E. coli* W with mature sequence numbers and structural features annotated. SS: periplasmic signal sequence; L: flexible linker; NT: unique N-terminal region; M60: peptidase/mucinase domain. **b,** Purified rSslE forms a clear ring of aggregated protein when incubated in citrate-phosphate buffer between pH 4.0 and 4.4 (upper panel; white asterisks). The ring formed above the liquid line is due to shaking of the sample and it resists solubilization up to pH 6.2 once formed (lower panel; purple asterisk), although some loss of protein band is observed after pH 5.8 (lower panel; red asterisks). **c,** Increase in ThT fluorescence emission on binding to rSslE fibres formed in citrate-phosphate buffer over pH 4.0 to 8.0. **d,** Immunoblot of rSslE incubated in citrate-phosphate buffer from pH 3.8 to 4.8 and detected with anti-His6 antibody. rSslE present in the well represents aggregated protein. **e,** Far-UV CD spectra of soluble rSslE at pH 4.4 (solid line) and 7.4 (dashed line). Secondary structure analysis is shown in the doughnut chart. **f,** ATR FT-MIR spectrum of the amide I region of rSslE fibres, displaying relatively high and low intensity absorption bands at ~ 1624 and 1695 cm^−1^ respectively, characteristic of an anti-parallel β-sheet structure.

Using electron microscopy (EM), small angle X-ray scattering (SAXS), nuclear magnetic resonance (NMR) spectroscopy and biochemical analyses we show that SslE from the *E. coli* W strain is formed of three defined regions (NT1, NT2 and NT3-M60), and is dynamic in solution. We also demonstrate that SslE undergoes conformational changes under acidic conditions, and this leads to the formation of higher order structures, which we propose is mediated by the NT2 and NT3 regions and is decorated with NT1 domains. We directly observe acidification within mature *E. coli* W biofilms and determine that SslE fibres can bind cellulose, a major exopolysaccharide of many *E. coli* biofilms. Furthermore, we provide evidence that SslE regulates the localization of exopolysaccharides during maturation of both *E. coli* W and ETEC strain H10407 biofilms and this is likely a general mechanism in all SslE-secreting bacteria.

## Results

### SslE forms aggregates with amyloid-like properties

Functional amyloids are important proteinaceous structures in biofilms^21^ and analysis of the SslE primary sequence had previously highlighted several regions across the protein that may form amyloid-like structures^22^. Furthermore, environmental pH plays a major role initiating the formation of amyloid-like structures in *Staphylococcus aureus*, *Enterococcus faecalis* and *Streptococcus mutans* biofilms^23, 24, 25^, and we speculated that pH may also stimulate similar structural changes in SslE. SslE from *E. coli* W, minus the N-terminal lipidation motif and adjacent disordered region (rSslE; residues 67 to 1497; UniProt ID E0IW31) (**Fig. 1a**), was therefore produced in *E. coli* K-12 and purified by nickel-affinity and size exclusion chromatography. We then incubated rSslE across the pH range 4.0 to 7.0 and observed a ring of protein deposited on the walls of the tube at pH ≤ 4.4, which resisted solubilization up to pH 6.2 (**Fig. 1b**). Mass spectrometry analysis of the ring identified peptides that spanned the complete sequence of rSslE and indicated that the aggregate contained intact protein, rather than a degradation product (**Supplementary Fig. 1**). We next used the amyloid diagnostic dye Thioflavin-T (ThT) to assess the formation of rSslE aggregates in solution over time. We observed large increases in ThT fluorescence emission again across the pH range 4.0 to 4.4, but no change was detected when rSslE was incubated above pH 4.4 (**Fig. 1c**). Likewise, when rSslE was boiled and treated with sodium dodecyl sulphate (SDS) and then analysed by immunoblotting, stable high molecular weight species were observed over the same pH range (**Fig. 1d**).

A main feature of amyloids is the presence of a cross-β-sheet structure^26^ and so we first analysed the secondary structure of soluble rSslE using far-UV circular dichroism (CD) spectroscopy. This suggested that at pH 7.4 rSslE is composed of approximately 30% α-helix, 12% β-sheet and 45% coil, while under acidic conditions and prior to aggregation, there is an ~8% loss of helical secondary structure with a ~5% increase in β-sheet structure (**Fig. 1e; Supplementary Table 1**). We next used attenuated total reflectance (ATR) Fourier transform mid-infrared (FT-MIR) spectroscopy to probe the secondary structure composition of soluble rSslE at pH 7.4 and rSslE fibres at pH 4.4. Examination of the amide I absorption band showed peaks at approximately 1656 cm^−1^ and 1624 cm^−1^, respectively (**Supplementary Fig. 2**). The shift in the amide I absorption band to a lower frequency indicated that a larger and more rigid structure had assembled under acid conditions. Furthermore, the presence of both a high-intensity absorption band at ~1624 cm^−1^ and a low-intensity band at ~1695 cm^−1^ in the pH 4.4 sample was indicative of an increase in anti-parallel β-sheet structure in rSslE aggregates^27^ and is consistent with previous FT-MIR based characterisations of other anti-parallel β-sheet amyloid fibres^28^ (**Fig. 1f**).

### Biofilm maturation supports polymerisation of SslE

Congo red is a dye which binds extracellular polymeric substances (EPS), including amyloids, and we used this to assess the role of SslE during the formation of macrocolony biofilms. We created an isogenic Δ*sslE* knockout mutant in *E. coli* W and ETEC strain H10407 and compared these to their parental wild-type strains. When these mutants were cultured on Congo red agar plates for 24 hrs, we observed no obvious differences, however, after 96 hrs the mutants exhibited an altered morphology with the dye retained within the centre of the colony (**Fig. 2a**). Complementation of the Δ*sslE* mutant in *E. coli* W with a plasmid containing intact *sslE* (Δ*sslE::sslE*) was able to restore bacterial secretion of SslE and recover macrocolony morphology similar to that of the parental wild-type (**Fig. 2a**; **Supplementary Fig. 3**). We also generated a plasmid expressing SslE with a truncated C-terminus (Δ*sslE::sslEΔM60*; residues 1 to 1000) to examine the role of the M60-domain during colony formation, and this again restored SslE secretion and wild-type colony morphology (**Fig. 2a**; **Supplementary Fig. 3**). This led us to speculate that SslE experiences extracellular pH values <5.0 during the development of biofilms and this results in SslE being deposited as a functional aggregate.

**Fig. 2.**
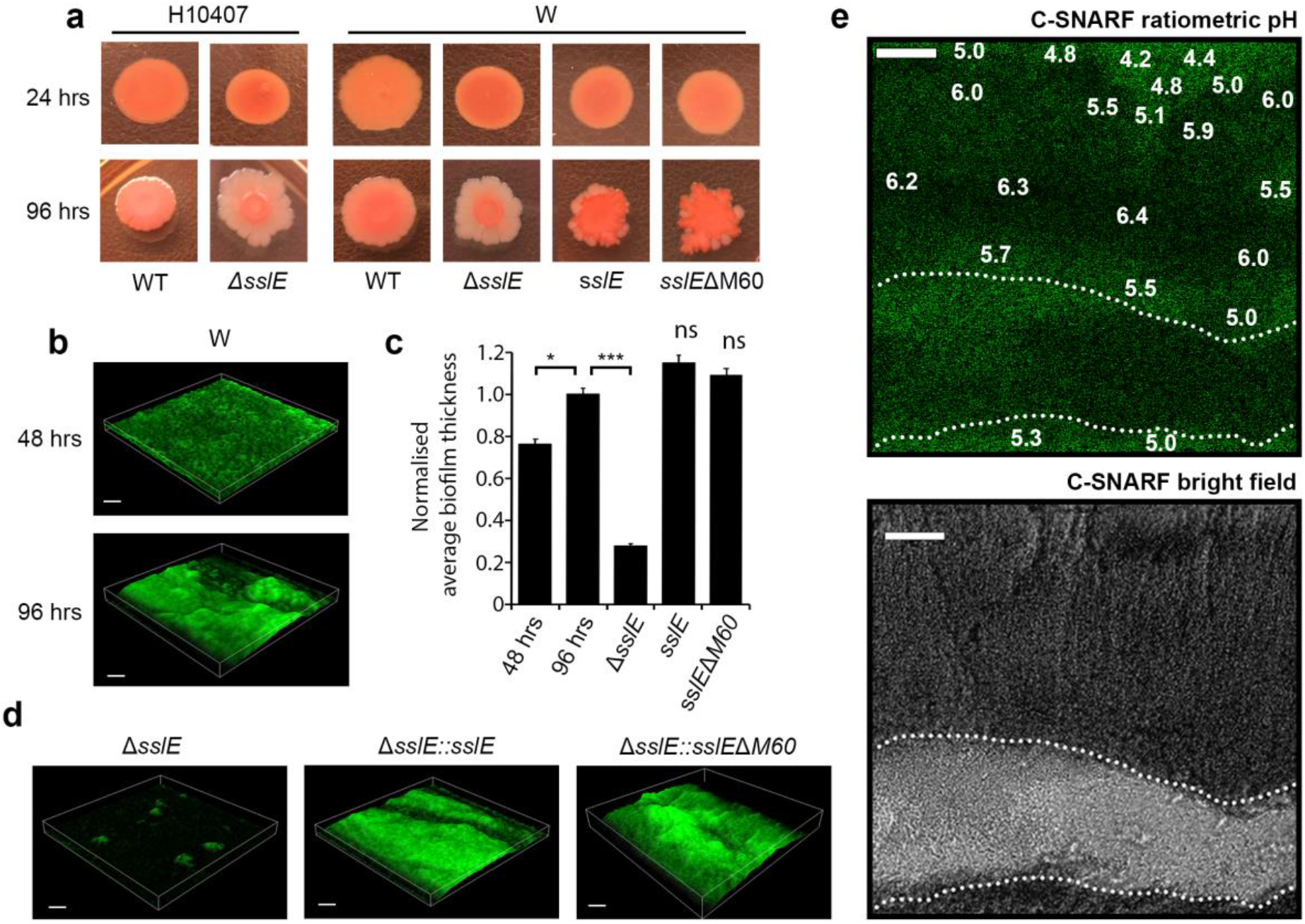
Analysis of SslE-dependent biofilm formation by *E. coli* W and H10407 strains. **a,** Macrocolony biofilm phenotype of wild-type *E. coli* W and H10407 strains and their derivatives on Congo red agar medium after 24 and 96 hrs of growth. The derivatives *sslE* and *sslE*ΔM60 represent *trans*-complementation of intact *sslE* or *sslE* minus its M60 domain, respectively, into the *E. coli* W Δ*sslE* mutant. **b,**CLSM images of wild-type *E. coli* W biofilms grown for 48 and 96 hrs and stained with lipophilic probe FM 1–43 (green). **c,** Quantification of 3-dimensional biofilm intensity of wild-type *E. coli* W biofilms and its derivatives carried out in ImageJ^40^. \**P* <.05; ^***^*P* <0.001; verses W 96 hrs growth by two-tailed Student’s t-test. **d,** CLSM images of *E. coli* W Δ*sslE* mutant and *trans*-complementation with *sslE* (Δ*sslE::sslE*) and *sslE*ΔM60 (Δ*sslE::sslE*ΔM60) stained with FM 1–43 (green). **e,** The pH across wild-type *E. coli* W biofilms grown for 96 hrs was monitored ratiometrically using C-SNARF-4 (green). pH values were calculated over ~30 μm^2^ boxes and pH values for representative regions within the biofilm fringes and centres are annotated. Dotted line outlines water channel within the biofilm, where pH values are not shown. Scale bar represents 20 μm. All data are representative of at least three independent experiments.

To test this, we first grew *E. coli* W biofilms using a microfluidic system under continuous flow and assessed their overall morphology using confocal laser scanning microscopy (CLSM). While biofilms grown for 48 hrs produced a relatively homogenous lawn of bacterial growth, after 96 hrs there were clear signs of maturation, with a significant increase in biofilm mass, structural heterogeneity and the presence of water channels (**Fig. 2b,c**). We then examined the *E. coli* W Δ*sslE* mutant under these conditions, but in line with previous reports in EPEC strain E2348/69^4^, it was unable to develop structures beyond microcolonies (**Fig. 2c,d**). Furthermore, complementation of the mutant with either intact *sslE* or *sslEΔM60* was again able to restore wild-type biofilm morphology. Together with our previous observation, this indicated that the C-terminal M60-domain is not required for translocation of SslE through its T2SS and it is dispensable for biofilm development, at least under these conditions.

We then examined pH distribution across *E. coli* W biofilms using CLSM coupled with the cell-impermeant fluorescent ratiometric probe seminaphthorhodafluor-4F 5-(and-6) carboxylic acid (C-SNARF-4). After growing *E. coli* W for 24 hrs under continuous flow, we observed pH values between 6.0 and 6.3 across biofilms (**Supplementary Fig. 4**), however, after 96 hrs, lower pH values were also recorded (**Fig. 2e**). Although across the majority of biofilms examined, we detected pH values between approximately 5.0 and 6.0, clearly defined microenvironments were also observed with pH values that ranged from 4.2 and 4.8.

We observed that when *E. coli* W was grown overnight in liquid culture at pH 5.0, the wild-type strain and both Δ*sslE::sslE* and Δ*sslE::sslEΔM60* complementation strains formed a clear ring on the wall of the tube that retained congo red dye (**Fig. 3a**). This was absent in the Δ*sslE* mutant or when grown at pH 7.0 and mass spectrometry analysis of the wild-type *E. coli* W ring again identified SslE peptides from across its sequence (**Supplementary Fig. 1**). Examination of wild-type *E. coli* W at pH 5.0 by immunoelectron microscopy revealed gold-labelled anti-rSslE antibodies associated with the bacterial outer membrane, but also associated with fibrous material in the extracellular milieu and emanating away from the bacterial surface; which was not observed in the Δ*sslE* mutant (**Fig. 3b,c,d**). Inspection of the *E. coli* W Δ*sslE::sslE* complemented strain under the same conditions produced less clustering of anti-rSslE antibodies, however, a wild-type phenotype was observed when cells were incubated at pH 4.0 prior to fixation and staining (**Fig. 3e,f**). This indicated that SslE can form aggregates upon secretion from *E. coli* and so we then used immunofluorescence to assess how SslE is distributed within established wild-type *E. coli* W and H10407 biofilms (**Fig. 4**). Here we observed a heterogenous distribution and clustering of SslE across biofilms formed by both strains, with SslE co-localised with bacteria, likely through association with the bacterial surface, but also clearly incorporated within the extracellular matrix.

**Fig. 3.**
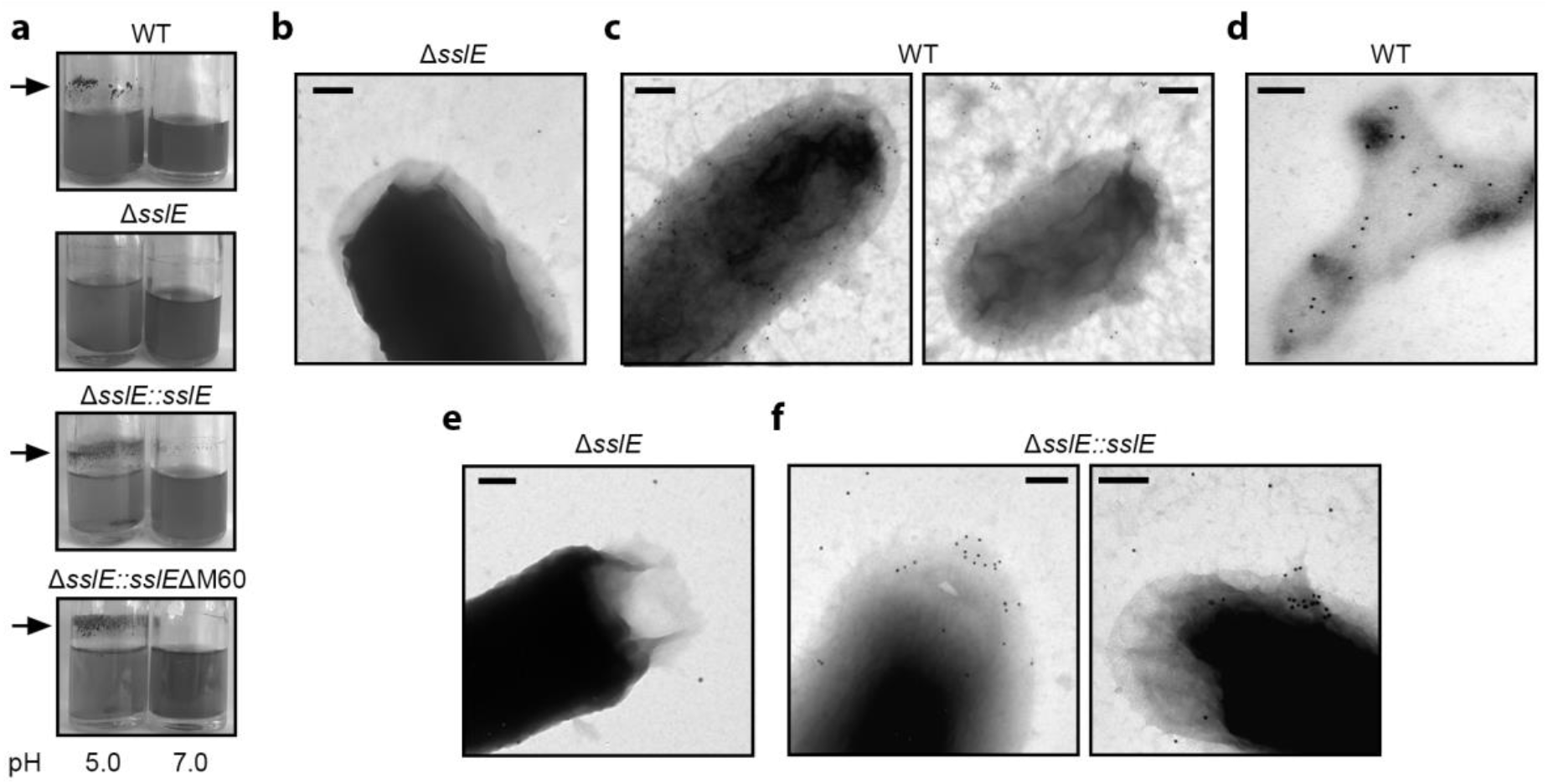
Analysis of SslE aggregation after bacterial secretion. **a,** Accumulation of SslE aggregates on the side of glass tubes containing overnight cultures of *E. coli* W strains, grown in LB media at pH 5.0 or 7.0. Arrows indicate Congo red stained ring containing SslE. **b-f,** SslE localization upon secretion from *E. coli* W strains assessed by immunoelectron microscopy. Bacteria were reacted with primary α-rSslE antibodies and secondary gold-labelled antibodies on carbon coated nickel grids and then negatively stained with uranyl acetate. **b, e,** *E. coli* W Δ*sslE* mutant washed in citrate-phosphate buffer at pH 5.0 or 4.0, respectively. **c,** Wild-type W strain washed in citrate-phosphate buffer at pH 5.0 with α-rSslE antibodies reacting with the bacterial surface and secreted fibrous material. **d,** Image highlighting clustering of antibodies to an extracellular aggregate. **f,** Δ*sslE::sslE* W strain washed in citrate-phosphate buffer at pH 4.0, again with α-rSslE antibodies reacting with the bacterial surface and secreted fibrous material. Scale bar is equivalent to 200 nm.

**Fig. 4.**
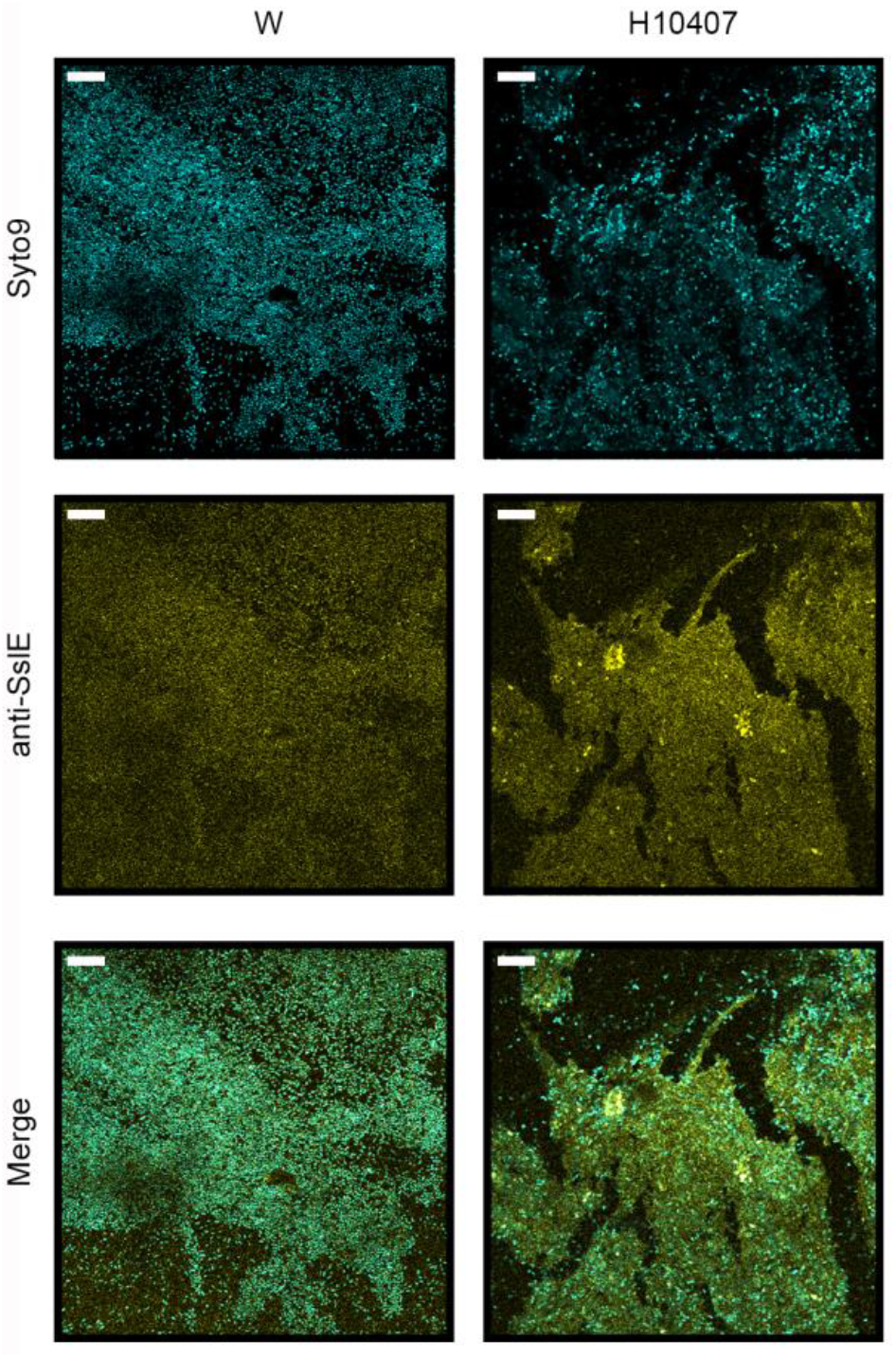
Localization of SslE within *E. coli* biofilms. Wild-type *E. coli* W and H10407 biofilms were formed using a continuous flow cell system over 96 hrs. Immunofluorescence were performed on fixed biofilms wherein the biofilms were incubated with α-rSslE antibodies (1:200 dilution) then incubate with goat anti-rabbit conjugated Alexa Fluor 633 (yellow). DNA was stained with Syto9 (cyan) to identify bacteria and eDNA within the biofilm matrix. Biofilms were visualized by CLSM. Scale bar is equivalent to 10 μm. All data are representative of three independent experiments.

### SslE aggregates bind cellulose

We speculated that during the maturation of biofilms, SslE aggregates interact with other components of the biofilm matrix and may contribute to the structural integrity of these communities. We therefore re-examined the morphology of wild-type and Δ*sslE* mutant *E. coli* W and H10407 macrocolonies using the carbohydrate stain calcofluor white, which binds β1-3 and β1-4 polysaccharides including cellulose (**Fig. 5a**). As with Congo red staining, calcofluor white extended to the edge of the colony in wild-type strains but was retained within the centre in the Δ*sslE* mutants. We then produced rSslE aggregates at pH 4.0 and isolated them in citrate-phosphate buffer at pH 6.0, a condition in which fibres would resist solubilization (**Fig. 1b**). As controls, we used monomeric rSslE, and the C-terminal domain (CTD) of the *Porphyromonas gingivalis* gingipain protein RgpB, both of which were not expected to bind cellulose or form polymers at this pH (**Supplementary Fig. 5**). We then assayed binding of rSslE fibres to immobilised cellulose discs and observed a significant interaction, but as expected this was not detected with monomeric rSslE or RgpB-CTD (**Fig. 5b**).

**Fig. 5.**
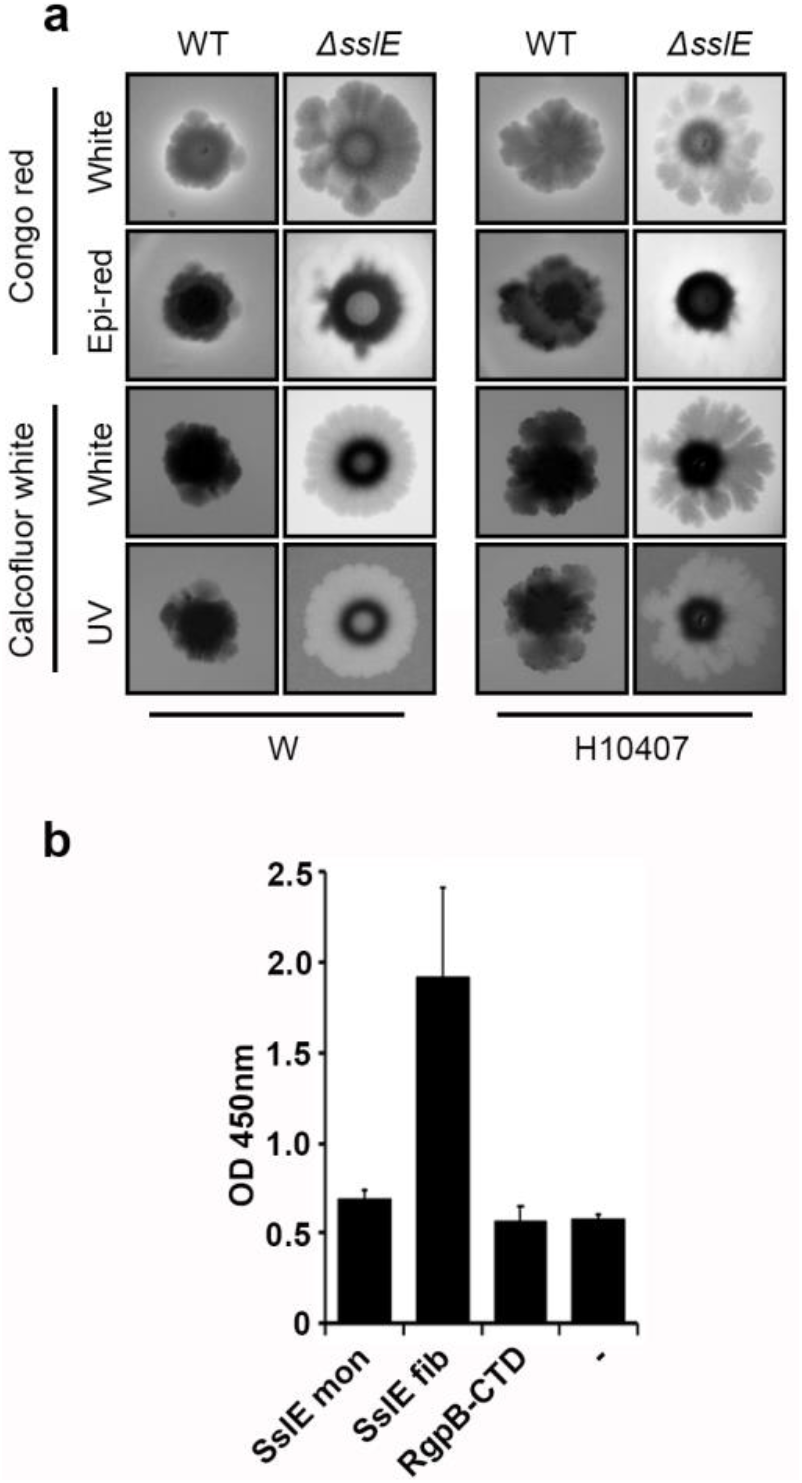
Binding properties of SslE aggregates. **a,** Macrocolony biofilm phenotype of wild-type and Δ*sslE* mutant *E. coli* W and H10407 strains on Congo red or calcofluor white agar medium after 96 hrs of growth. Macrocolonies were visualized with white light to determine the edge of the colony, and then either epi-red or UV light was used to determine the localization of Congo red or calcofluor white, respectively, within the colony. **b,** ELISA analysis of binding between cellulose discs and His-tagged rSslE monomers, His-tagged rSslE fibres and His-tagged *P. gingivalis* RgpB-CTD control, at pH 6.0 and detected with anti-His6 antibody. BSA-coated wells were used as controls. \**P* <0.05; verses control empty well (-) by two-tailed Student’s test. All data are representative of at least three independent experiments performed in triplicate.

### Overall structural features of SslE

We next initiated structural studies of SslE. Small angle X-ray scattering (SAXS) coupled with size exclusion chromatography (SEC-SAXS) was first used to get shape information for rSslE at pH 7.4. Guinier analysis provided a radius of gyration (*R*_g_) of 4.03 nm and examination of the distance distribution function (*P(r)*) suggested a maximum particle dimension (*D*_max_) of 14.1 nm and *R*_g_ of 4.07 nm (**Supplementary Fig. 6; Supplementary Table 2**). Calculation of the particle molecular mass (151 kDa) was consistent with a monomer in solution (theoretical mass 160 kDa). *Ab initio* dummy residue reconstruction generated 20 reproducible models with an average normalized spatial discrepancy (NSD) score between reconstructions of 0.56 and a χ^2^ fit between calculated and experimental solution scattering of 1.2 (**Fig. 6a,b**). The averaged low-resolution bead model suggested that rSslE has a slightly elongated structure, however, examination of the Kratky plot indicated that rSslE is a dynamic particle (**Supplementary Fig. 6**) and no obvious domain features could be assigned.

**Fig. 6.**
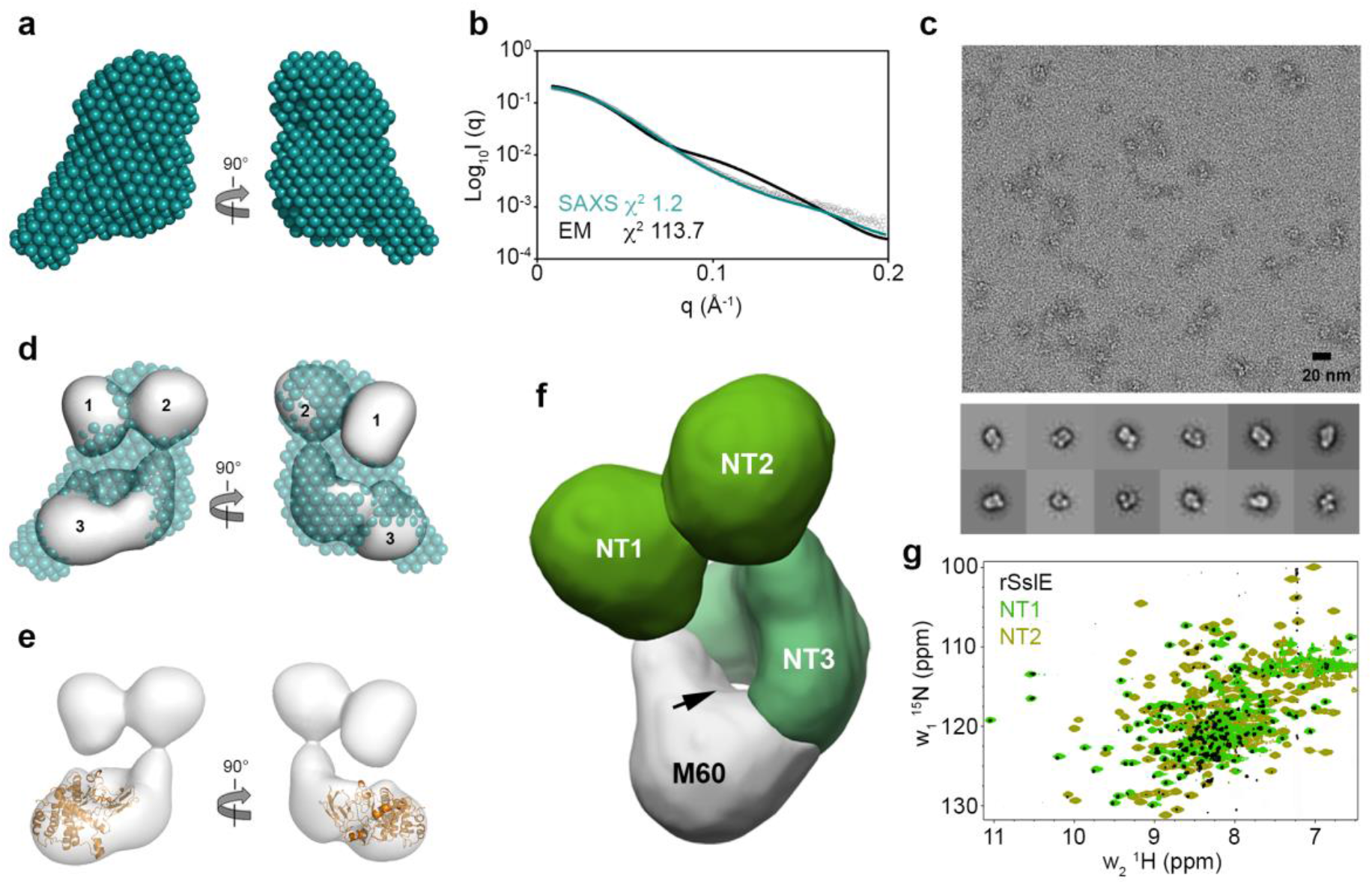
Structural features of monomeric SslE. **a,** SAXS bead model of rSslE at pH 7.4. **b,** Fit of the SAXS bead model (teal line) and negative-stain TEM map (black line) of rSslE to the rSslE SAXS data (black open circles) with χ^2^ of 1.2 and 113.7, respectively. **c,** Representative negative-stain TEM micrograph of rSslE (pH 7.4) at 50,000× nominal magnification with representative 2D classifications. Scale bar represents 20 nm. **d,**Overlay of the 22 Å resolution TEM map (grey) and SAXS bead model (teal) with the three defined regions in SslE highlighted. **e,** Docking of an SslE M60 domain homology model (orange) into the TEM map. **f,** TEM map of rSslE colored based on domain organization. The NT3-M60 interdomain channel is highlighted with an arrow. **g,** TROSY ^1^H^15^N-HSQC spectrum of rSslE (black) overlaid with ^1^H^15^N-HSQC spectra of the SslE NT1 (light green) and NT2 (olive green) domains.

Negative-stain TEM data was next collected for rSslE at pH 7.4 and we performed single particle analysis to generate an initial low-resolution structure, with overall dimensions of approximately 8×8×11.5 nm (**Fig. 6c; Supplementary Fig. 7**). Here, three well defined regions could be identified: two adjacent globular domains (regions 1 and 2) with approximate diameters of 4 nm and 4.5 nm, respectively, and a torus shaped structure (region 3) with a ~15 Å wide central channel and approximate overall dimensions of 4.5×6.5×9 nm (**Fig. 6d**). Superposition of the SAXS bead model with the TEM envelope supported SslE being dynamic in solution, with an apparent reduction of bead volume in all three regions of the SAXS model. Docking analysis of a homology model of the M60 domain, based on the IMPa M60 metalloprotease from *Pseudomonas aeruginosa* (PDB ID code 5kdv)^19^, indicated that the C-terminus of SslE is located at the base of region 3 (**Fig. 6e**). This suggested that the remaining density in region 3 is composed of the SslE sequence directly N-terminal to the M60 domain and these two regions pack against one another to form a stable structure around an inter-domain channel. Furthermore, from this model the M60 HExxH active site motif faces into the channel and this could be an important feature for its mucolytic activity. The location of the M60 domain in the TEM map was also supported by previous reports where it was not possible to purify a folded recombinant SslE M60 fragment due to sample instability^22^.

Secondary structure analysis^29^ of the SslE sequence indicated two potential inter-domain boundaries within the N-terminal region and we predicted that these represented regions 1 and 2 located at the extreme N-terminus (NT) of SslE. We therefore renamed these as the NT1 (residues 67-211) and NT2 (residues 230-425) domains, respectively, and renamed the remaining sequence as NT3 (residues 426-1000) (**Fig. 6f**). The NT1 and NT2 domains, along with the NT1-NT2 region (residues 67-425) and NT3-M60 core (region 3; residues 426-1497) were then produced in *E. coli* K-12 and purified by nickel-affinity and size exclusion chromatography. Examination of these constructs using analytical gel filtration and nuclear magnetic resonance (NMR) spectroscopy showed that they were well folded (**Supplementary Fig. 8**), and supported NT1, NT2 and NT3-M60 being well-defined structural boundaries within SslE. When we examined intact rSslE by NMR, a ^1^H^15^N transverse relaxation optimized spectroscopy (TROSY) HSQC spectrum yielded only ~250 strong resonances with the remaining ~1200 peaks being either very weak or completely absent (**Fig. 6g**). Furthermore, NMR relaxation measurements of these intense peaks provided an estimated correlation time consistent with an ~80 kDa domain (τ_c_ ~ 35 ns at 37°C). Comparison of rSslE with ^1^H^15^N-HSQC spectra from SslE subdomains also clearly showed that the rSslE NMR spectrum is dominated by the NT1 domain but with minor contributions from the NT2 domain (**Fig. 6g; Supplementary Fig. 9**). This further supported this region having significant independent motion with respect to the NT3-M60 core and we considered whether the NT1-NT2 region alone could be responsible for mediating SslE aggregation. We therefore repeated our ring assay with SslE NT1-NT2 and the NT3-M60 constructs incubated across the pH range 4.0 to 7.0. This resulted in significantly less protein being deposited on the walls of the tubes and for NT3-M60 this was over a wider pH range (**Supplementary Fig. 10**). This implied that both the NT1-NT2 and the NT3 regions are necessary for correct polymerisation of intact SslE.

### Kinetics of SslE aggregation

We took advantage of SEC-SAXS to probe the global shape of rSslE at pH 4.4 prior to fibre formation and both Guinier analysis (*R*_*g*_ 3.92 nm) and examination of the distance distribution function (*D*_*max*_ 13.7 nm) produced very similar values to rSslE at pH 7.4 (**Supplementary Fig. 11; Supplementary Table 2)**. However, the presence of better-defined features in the SAXS profile at acidic pH and the differences at large q-values in the normalized Kratky plots were indicative of rSslE becoming more rigid under acidic conditions (**Fig. 7a**; **Supplementary Fig. 12**). *Ab initio* dummy residue reconstruction produced 20 models with an average NSD score of 0.45 and a χ^2^ fit between calculated and experimental solution scattering of 1.0 (**Fig. 7b**; **Supplementary Fig. 13**). Comparison of the averaged bead models indicated an increase in bead volume within the NT1 and NT2 domains and adjacent NT3 region at pH 4.4, while little change was observed in the NT3-M60 core, which supported an increase in rigidity under acidic conditions that could trigger aggregation.

**Fig. 7.**
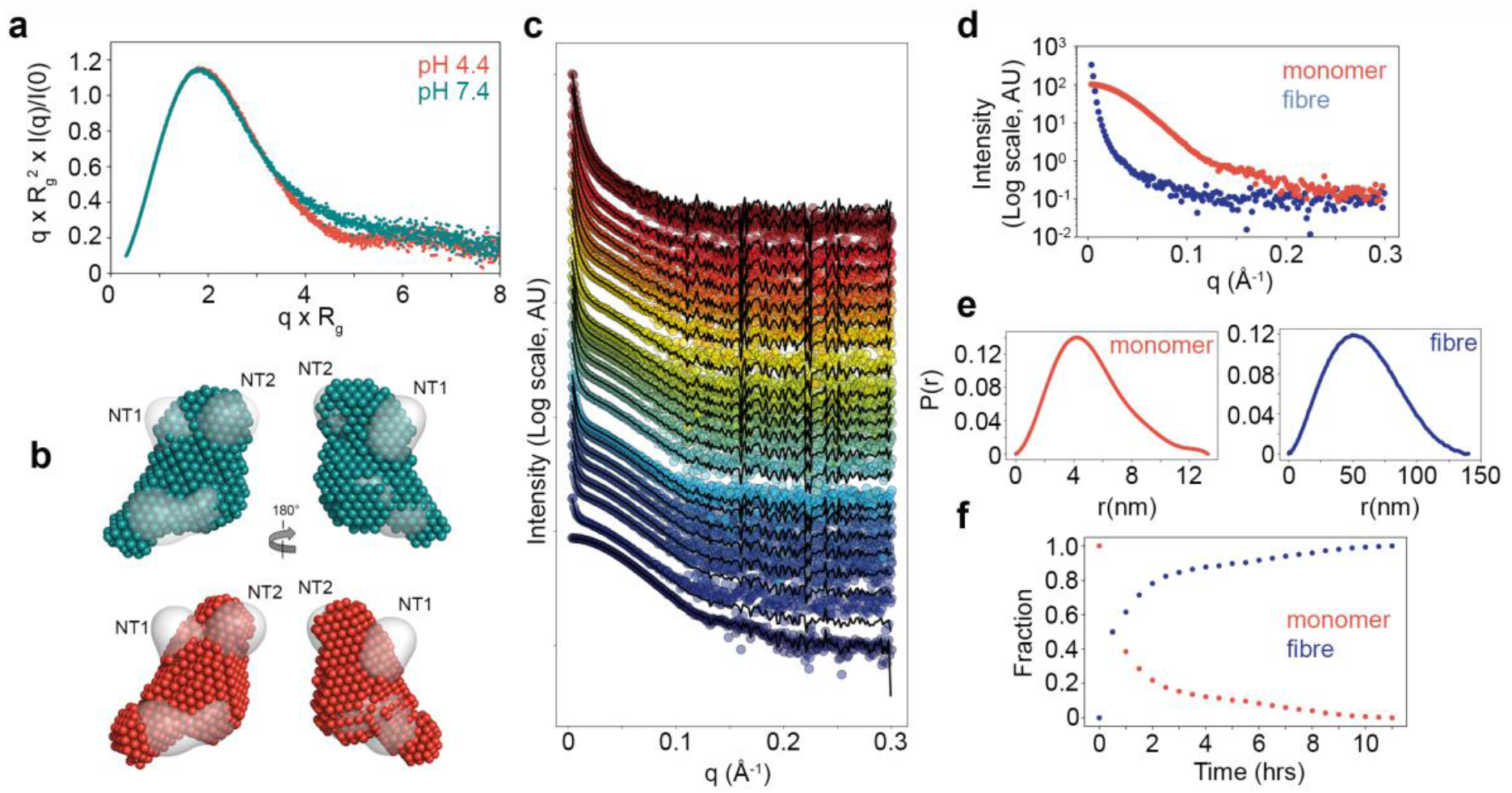
Kinetics of SslE aggregation. **a,** SAXS normalized Kratky plots for rSslE at pH 4.4 and 7.4. **b,** Overlay of rSslE SAXS bead models at pH 4.4 (red) and 7.4 (teal) with rSslE TEM map (grey). NT1 and NT2 domains are highlighted. **c,** COSMiCS decomposition of rSslE SAXS fibrillation data, assuming the co-existence of two components. SEC-SAXS curve for rSslE at pH 4.4 is also shown for t=0. **d,** COSMiCS decomposed scattering curves for rSslE monomer (red) and aggregates (blue). **e,** Shape distribution [*P(r)*] functions derived from COSMiCS decomposed scattering curves for rSslE monomer (red) and aggregates (blue). **f,** Relative populations of the two species of rSslE derived from the COSMiCS analysis over time.

We then exploited the additive nature of SAXS along with its ability to examine structures over a large size range to study the kinetics of fibre formation by rSslE. We acquired SAXS data at pH 4.4 during the course of aggregation for 11 hrs and the resulting 23 profiles displayed a systematic upwards intensity increase, indicating the presence of large aggregates in solution (**Fig. 7c**). The appearance of these large species occurred relatively quickly as the upwards intensity increase was already observed after 1 hr and the increase of aggregated species was concomitant to the loss of the SAXS features linked to the monomeric species. Principal Component Analysis (PCA) indicated that the data set could be described as a two-component system (**Supplementary Fig. 14**) and we decomposed the time-dependent data set along with the SAXS profile of the monomer, obtained using SEC-SAXS, with COSMiCS. COSMiCS uses a chemometric approach to decompose SAXS data sets in a model-free manner to derive the pure SAXS curves of the co-existing species and their relative populations, which we used here to report on the fibrillation kinetics^30^. The COSMiCS decomposition, assuming the co-existence of two components, was able to adequately fit all the SAXS profiles with an average <χ^2^> of 1.5, although some deviations from the perfect fit were observed for the SAXS curves measured in the first 3 hrs (**Fig. 7c**). This observation suggested the presence of a small population of a third species in the initial steps of fibrillation but this putative third species could not be captured when considering a three-species decomposition with COSMiCS.

The extracted profile of the smallest species, and the subsequently derived *R*_g_ (3.93 nm) and *D*_*max*_ (13.3 nm) values, were almost identical to the SEC-SAXS profile for rSslE at pH 4.4 (**Fig. 7d,e; Supplementary Table 2**), and this indicated that SslE is in a monomeric form at the beginning of the fibrillation process. The extracted SAXS profile of the second component displayed the typical features of a large particle and *P(r)* analysis indicated it had an *R*_*g*_ of ~51.1 nm and *D*_*max*_ of ~140 nm (**Fig. 7d,e; Supplementary Table 2**). Fractal fit analysis^31, 32^ of the decomposed SAXS profiles also indicated that rSslE aggregates into a structure that is fractal in nature (**Supplementary Fig. 15**). The radius of the primary particle was determined to be ~3.5 nm while the mass fractal and surface fractal dimensions were determined to be 2.9 and 3.0, respectively. Examination of the relative populations of the two species derived from the COSMiCS decomposition over time showed that after 30 min from initiating the measurements, the aggregated species already represented 50% of the molar fraction (**Fig. 7f**). From this time point, this population continued to grow and reached a plateau after ~10 hrs.

### Structural model of SslE fibres

Negative-stain TEM was used to visualize purified rSslE fibres revealing two common morphologies (**Fig. 8a**). The first form appeared as short single, flexible fibres with a core structure measuring ~4.5 nm wide by ~110 nm in length and decorated with globular structures ~4.5 nm in diameter (**Fig. 8b,c**). The second morphology again resembled a fibrous material but measured between ~20 to 40 nm in width by 200 to 300 nm in length (**Fig. 8d**) and appeared to be an aggregation of the smaller fibres. Further analysis of rSslE aggregates in solution by real-time multi-angle light scattering (RT-MALS) supported these observations (**Supplementary Fig. 16**). We measured a smaller species with an average molecular mass of 5.4×10^3^ kDa (± 0.4×10^3^) and a larger more polydisperse species with an average molecular mass of 12×10^3^ kDa (± 0.8×10^3^). In addition, we measured an average *R*_g_ of 137.5 nm (± 1.7) and 139.8 nm (± 1.5) for these species, respectively, which is in line with *R*_g_ measurements using COSMiCS. Examination of rSslE fibres using solid state NMR (ssNMR) showed individual narrow signals (**Fig. 8e,f**), indicative of an ordered structure, and we could tentatively assign approximately 40 ordered residues to specific residue types. Although we could not assign these resonances to a specific sequence position in SslE, the pattern suggested that these residues were localised to more than one site in SslE.

**Fig. 8.**
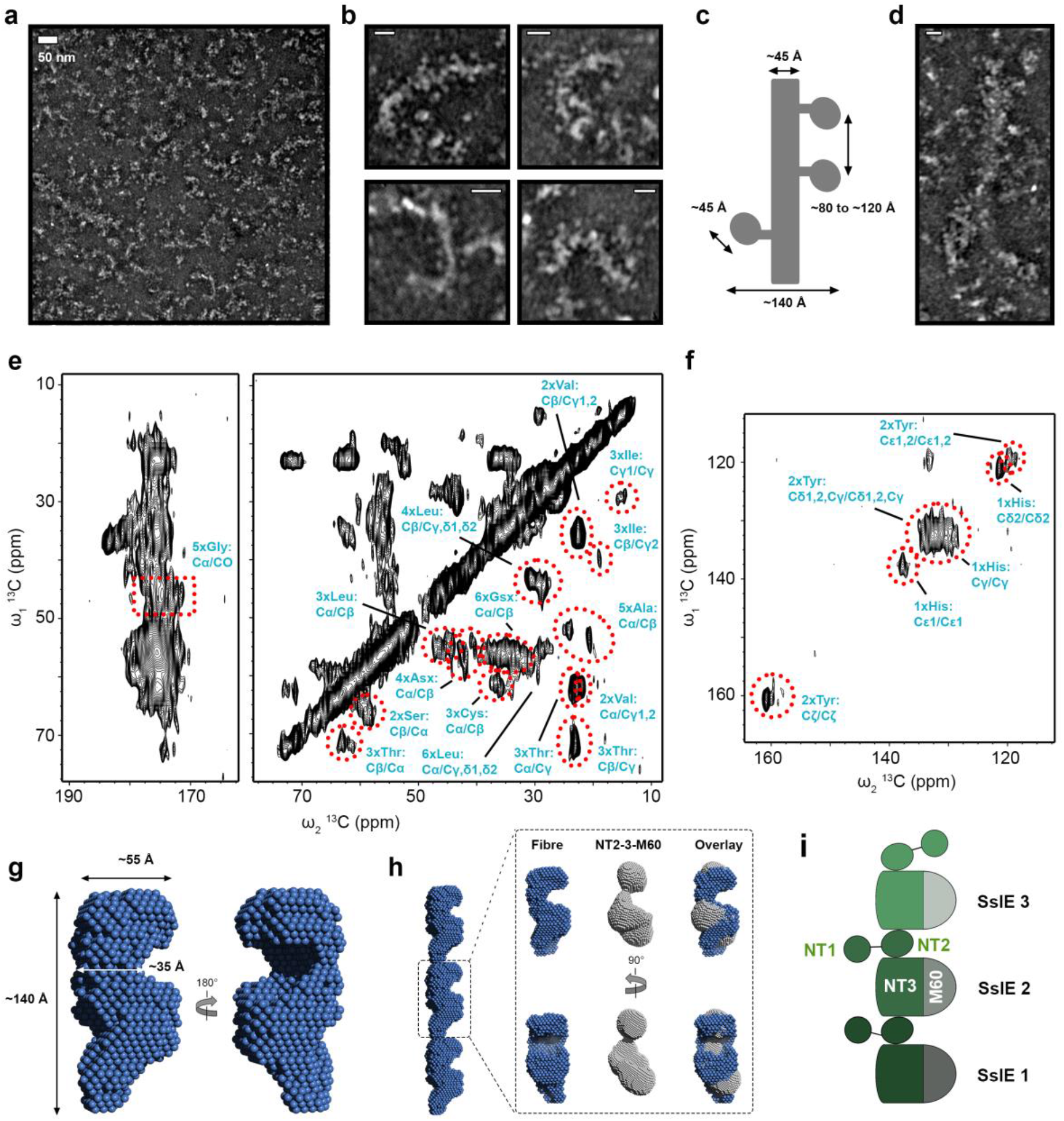
Model of SslE fibres. **a,** TEM analysis of negatively stained SslE fibres. Two major species are observed. **b,**Smaller filaments have an average width of ~4.5 nm and are coated with globular structures of ~4.5 nm in diameter. Scale bar, 20 nm. **c,** Schematic describing the overall dimensions of the smaller SslE fibre species. **d,** Larger species of SslE fibres appear as ~20 to 40 nm wide structures with variable lengths. Scale bar, 20 nm. **e, f,** DARR ssNMR spectrum of SslE fibres focussed on the carbonyl (**e,**left), aliphatic (**e,** right) and aromatic (**f**) regions. Likely amino acid type assignments are highlighted with red dashed lines and annotated in teal. **g,** S AXS bead model for the repeating unit of the SslE fibre core with overall dimensions given. **h,** Three units of the SslE fibre core translated along the fibre axis (left). One unit is zoomed in and overlaid with a bead model created of the rSslE negative-stain TEM map with the NT1 domain removed (NT2-3-M60; right). **i,** Proposed model for SslE fibres. The core of the fibre contains the NT2, NT3 and M60 domains and is decorated with flexible NT1. The core fibre is formed via interactions between the NT2 and NT3 domains of adjacent SslE molecules.

As the SAXS fibrillation data at the 10 hr time point had shown depletion of monomeric rSslE but also had reduced fractal properties compared with later time points (**Fig. 7f**), we discarded the low q region and used this data to carry out analysis of the fibre asymmetric unit. Geometrical shape analysis suggested that rSslE fibres form a cylinder with radius 3.2 nm and height 16.5 nm, and cross-sectional analysis indicated a repetitive unit with a cross-sectional *R*_*g*_ (*R_c_*) of 4.78 nm. The cross-sectional *P*(*r*)*C* gave a cross-sectional *D*_*max*_ of 14.0 nm. *Ab initio* dummy residue reconstruction produced a final model with a χ^2^ fit between calculated and experimental solution scattering of 1.6 (**Fig. 8g; Supplementary Fig. 17**). The width of the repeating unit varied between ~3.5 and ~5.5 nm, which is consistent with our measurements of single fibres by TEM. As we could not identify peripheral globular domains in the fibre bead model, this supported SslE fibres being formed from a central core structure, with folded and dynamic domains presented on the surface (**Fig. 8h**). Comparison of the fibre reconstruction with bead models derived from the rSslE TEM envelope also suggested that a single molecule of SslE would satisfy the bead volume but only once the NT1 domain had been removed.

## Discussion

In this study we have revealed that SslE is composed of four defined regions: two globular domains at the N-terminus (NT1 and NT2) and a central torus shaped core, where a NT3 domain packs against an M60-like aminopeptidase. The N-terminal region (NT1, NT2 and NT3) is composed of a unique primary sequence that has only been identified in SslE and its homologs^33^, while the C-terminal M60-like domain is found in a wide range of proteins secreted by both prokaryotic and eukaryotic microorganisms that interact with mucosal membranes^18, 34^. The atomic structures for three M60-like domains in complex with *O*-glycopeptides have been described: BT4244 from *Bacteroides thetaiotaomicron* (PDB ID code: 5kd2), IMPa from *P. aeruginosa* (PDB ID code: 5kdw) and ZmpB from *Clostridium perfringens* (PDB ID code: 5kdn)^19^. While the active site structures in these bacterial enzymes are highly conserved, there are variations within the remaining M60 domain, which enables them to recognize unique glycan sequences. Structural data is not available for intact BT4244 or ZmpB, although they are thought to contain additional carbohydrate binding domains^18^ and it is likely that these M60 augmentations have a role in the recognition of *O*-glycopeptide substrates. It also appears that the active site in the M60 domain of SslE faces into the NT3 domain and this implies that both the NT3 and M60 domains are responsible for the recognition of mucins in SslE. With significant flexibility of the NT1 and NT2 region, these domains may also interact with mucin substrates and may promote processivity.

Although SslE has a clear function in the processing of mucins, it is a unique member of the M60-like aminopeptidase family as it is also fundamental for the maturation of EPEC biofilms grown on plastics^4^. Furthermore, we have now shown that SslE is also required for the maturation of *E. coli* W and ETEC H10407 biofilms when grown statically on agar and under shear stress on glass substrates, where it is widely distributed throughout the extracellular matrix. We have demonstrated that this unusual function is due to SslE forming aggregates at pH ≤ 4.4; conditions which are also observed as microenvironments across mature wild-type *E. coli* W biofilms. It is well established that phenotypical heterogeneity develops during the maturation of biofilms and direct measurement of extracellular pH within *E. coli* PHL628 biofilms has also shown a heterogenous distributions of pH, with values ranging between 5.0 and 7.0^35^. Therefore, it is very likely that SslE forms functional aggregates within transiently forming microenvironments across maturing and established biofilms, but then resists solubilisation after any increase in the local pH.

Microbial amyloid structures are ubiquitous in biofilms and in gram negative bacteria most known amyloid structures are synthesized by either the Curli or Fap secretion systems^36, 37, 38, 39^. These form through spontaneous aggregation of extended peptides and provide adhesion, cohesion and contribute to the structural integrity of the biofilm matrix. Although our biochemical and biophysical analyses suggests that under acidic condition rSslE displays some amyloid-like features, examination of aggregates using negative-stain TEM has revealed short fibres associated with globular structures. SAXS reconstructions of rSslE fibres has allowed additional analysis and suggests that the repetitive fibre unit can accommodate a single NT2-NT3-M60 region, and we therefore propose a potential model for the fibrillation of SslE into functional aggregates via end-to-end stacking (**Fig. 8i**). In this model, the asymmetric unit of the fibre core is composed of the NT2, NT3 and M60 domains and the NT1 domain is flexible and can move freely around the surface. Polymerisation of the fibre core is mediated through ~40 residues within the NT2 and NT3 domains but not the M60 domain, but this would require some rearrangement of the NT3/M60 interface.

FTIR analysis of rSslE aggregates provided an amide I maxima of 1624 cm^−1^, which is consistent with an increase in anti-parallel β-sheet conformation and also lies within the range of amide I maxima for other typical amyloid structures (*e.g.* 1611-1630 cm^−1^)^40^. However, larger and more rigid amyloid structures generally have peaks at lower frequencies, while native β-sheets show absorption bands with maxima between 1630-1643 cm^−1^. This suggests that the rSslE amide I band peak represents an amyloid structure which is relatively small and is not overly rigid, although it could alternatively originate from fibre stacking/clumping effects. Binding of ThT to rSslE at acidic pH also supports a general increase in β-sheet content in fibres but could also indicate the formation of new cavities upon fibrillation^41^. It is therefore still unclear whether these changes reflect the formation of a continuous amyloid-like structure that runs the length of the fibre core or instead represent β-sheet augmentation between adjacent SslE molecules along the fibre, upon fibrillation of SslE.

We have shown that SslE influences the localization of exopolysaccharides during the development of macrocolony biofilms and SslE fibres can directly associate with cellulose, which is known to have a structural function in many *E. coli* biofilms and provides cohesion and elasticity^42^. Although the precise role of SslE during biofilm development is still vague, it could have an architectural role acting as a hub to promote binding between other EPS or it could regulate the correct deposition of EPS within the biofilm. As monomeric SslE does not recognise cellulose, this suggests that new binding sites are formed after its aggregation. The presence of a surface exposed domain also implies that this domain may be functional in binding cellulose or other yet to be identified ligands, or is functional under different conditions, for example through interacting with mucins during intestinal colonisation. Likewise, the fibre core may still retain its mucolytic activity and could play a role in mucin restructuring during biofilm growth in the gut.

Although both *E. coli* W and ETEC 10407 secrete SslE, the W strain is a harmless commensal, while ETEC is a major etiological agent in developing countries^43^. SslE is also required for colonisation in a rabbit model of EPEC infection^4^ and is actively transcribed during ETEC infection of mice^6^. Biofilm growth and the production of enterotoxins are essential for the pathogenesis of ETEC-induced disease^43^ and secretion of SslE and the ETEC labile toxin is via the same type II secretion system^4, 44^. This suggests that SslE increases the virulence of *E. coli* pathotypes through its ability to promote biofilm maturation and/or through its interactions with mucosal defences. Both enterotoxigenic and enteropathogenic *E. coli* strains cause infection in the small intestine^1^ where the intraluminal pH ranges between approximately 6.0 to 7.5^45^. However, *E. coli* possess an acid-tolerance response that supports exponential growth at pH values as low as pH 4.2^46^ and due to the synthesis of organic acids by the residing microbiota, during colonisation of the intestine *E. coli* will experience extracellular pH values that range between 4.0 and 6.0^47^. Although this suggests that within the intestinal tract, conditions exist where SslE could regulate biofilm development, it remains unclear whether this role is important for infection or during extraintestinal survival. SslE represents a unique protein and further studies are now required to understand its complete functions during ecology and disease.

## Methods

### Bacterial strains and media

All bacterial strains, plasmids and primers used in this study are listed in **Supplementary Table 3**. Lysogeny broth (LB) medium contained (per litre) 10 g tryptone, 5 g yeast extract and 10 g NaCl. Isotopically defined M9 minimal medium (pH 7.4) contained (per litre) 6.0 g Na_2_HPO_4_·7H_2_O, 3 g KH_2_PO_4_, 0.5 g NaCl, 0.12 g MgSO_4_·7H_2_O, 0.022 g CaCl_2_, 0.04 g thiamine, 8.3 mg FeCl_3_·6H_2_O, 0.5 mg ZnCl_2_, 0.1 mg CuCl_2_, 0.1 mg CoCl_2_·6H_2_O, 0.1 mg H_3_BO_3_ and 13.5 mg MnCl_2_·6H_2_O, supplemented with 2g [U-^13^C_6_]glucose and/or 0.7 g ^15^NH_4_Cl. M9 media was made up in deuterium oxide for the production of perdeuterated protein samples and pH was adjusted using 1 M NaOH solution. All NMR isotopes were from Sigma.

### Gene deletion

A non-polar deletion of *sslE* was constructed in *E. coli* W and H10407 strains by allelic exchange with FLP recombination target (FRT)-flanked Kan^r^, using a modified protocol^48^. This fragment was amplified from *E. coli* strain JW5925-1^49^ using Platinum *Taq* DNA polymerase (Promega) and primers PC3 and PC4 with specificity for the regions flanking *sslE*. These exchanges were facilitated by the λ Red recombinase system carried on plasmid pKD46 and knockouts were confirmed by PCR and sequencing.

### Plasmid construction

Complementation plasmids pCPC1 and pCPC2 were generated by amplifying whole length *sslE*, or *sslE* minus the M60 domain, from *E. coli* W gDNA using primer pairs PC7/PC8 or PC7/PC9, respectively. These were digested with HindIII/NheI (NEB), ligated into HindIII/NheI-digested pBad-cm18 vector. Synthetic genes gPC4, gPC5, gPC6 and gPC7 were synthesized by Synbio Technologies and cloned into pET28b vector using NcoI and XhoI restriction sites to create plasmids pPC2, pPC3, pPC4 and pPC5, respectively (**Supplementary Table 4**). Plasmid pBD1 was created by amplification of the RgpB-CTD from *P. gingivalis* W50 gDNA using primers BD1 and BD2. This was then cloned into pET46 Ek/LIC vector through ligation independent cloning (Novagen).

### Microfluidic biofilm growth

Biofilms were grown in a BioFlux 200 device (Labtech, UK) using a protocol adapted from the manufacturer. Single colonies of *E. coli* W and its derivatives were resuspended in 10 ml of LB (with additional 50 μg/ml kanamycin for *Δssle* mutant and 50 μg/ml kanamycin, 25 μg/ml chloramphenicol *Δssle::pCPC1* (*sslE*) and *Δssle::pCPC2* (*sslEΔM60*)) and incubated at 37°C with shaking (200 rpm) for 16 hrs. All subsequent media contained appropriate antibiotics where required. The cultures were then diluted (1:100) in 10 ml of LB to OD_600nm_ of 0.5. A 24 well Bioflux plate (0-20 dyn/cm^2^) was pre-warmed to 37°C on a heated stage and microfluidic channels were incubated with prewarmed LB. The channels were then inoculated by injecting 20 μl of bacterial suspension into the output reservoir for 5 s at 2 dyne/cm^2^. The microfluidic plate was incubated for 1 hr at 37°C to allow bacteria to bind to the surface and then the flow reversed for 20 s. Prewarmed LB was then added to the input reservoir, the flow of media was initiated at 2 dyne/cm^2^ for 5 min and then decreased to 1 dyne/cm^2^ for 18 hrs. The spent media was removed from the output reservoir, fresh prewarmed media was added to the input reservoir and the flow was lowered to 0.5 dyne/cm^2^ for up to 96 hrs.

### Biofilm pH measurement

The ratiometric dye seminaphthorhodafluor-4F 5-(and-6) carboxylic acid (C-SNARF-4) was used to directly measure pH across biofilms within the Bioflux channel, using a modified method^50^. HEPES buffer was first adjusted between pH 3.2 to 7.8 in 0.2 increments using HCl, and 95 μl of each respective solution was then mixed with 5 μl 1mM C-SNARF-4, resulting in a final 50 μM concentration of C-SNARF-4. This solution was flushed into the Bioflux plate channel, pre-warmed at 37°C and the channels were imaged at x64 magnification using a DM-IRE2 confocal laser scanning microscope (Leica Microsystems Heidelberg GmbH, Germany) and the accompanying Leica Microsystems Confocal Software (version 2.61 Build 1537). Simultaneous images were captured using an excitation wavelength of 488 nm and emission wavelengths of 580 nm and 640 nm, in triplicate for each pH increment with a background image obtained after each acquisition. Acquired images were analysed using ImageJ Software (FIJI)^51^ by determining the average intensity of fluorescence and standard deviation of each channel, minus the background control. The intensity ratio for each pH was used to make a standard curve by plotting known pH against green/red ratio. 50μM C-SNARF-4 was then flushed into Bioflux plates containing *E. coli* W biofilm after either 24 or 96 hrs growth and five separate images were acquired for each Bioflux channel to calculate the pH from different areas of the biofilm. The fluorescence intensity and standard deviation were recorded, and the ratiometric value was used to determine pH from the standard curve. A 12×12 grid (~30 μm^2^ area) was applied to each image; each box was used to define regions of interest (ROI) and an average measure of pH was calculated within each region. The average pH for ROIs within the biofilm fringes and centres is presented.

### Biofilm immunofluorescence

Single colonies of wild-type *E. coli* W or H10407 were resuspended in 10 ml of LB and incubated at 37°C with shaking (200 rpm) for 16 hrs. Cultures were diluted (1:100) in prewarmed LB and 5 ml was injected onto borosilicate glass coupons held within FC310 flow cells (Biosurface Technology) and then incubated for 1 hr. Flow of LB media was initiated at 0.4 ml/min and maintained for 96 hrs. Coupons were removed and washed three times for 10 min each in PBS-Tween. Coupons were then blocked for 1 hr in 2% (w/v) BSA, PBS-Tween at room temperature and incubated overnight at 4°C with polyclonal anti-rSslE antibody (rabbit; Invitrogen), diluted 1:200 in 0.1% (w/v) BSA, PBS-Tween. After three 10 min washes in PBS-Tween, coupons were incubated in the dark for 1 hr with 10 μM Syto9 (Invitrogen) and anti-rabbit Alexa Fluor 633 secondary antibody (goat; Invitrogen) diluted 1:500 in 0.1% (w/v) BSA, PBS-Tween. This was followed by three 10 min washes in the dark in PBS-Tween and then fixation for 15 min at room temperature in the dark with 3% (w/v) paraformaldehyde, PBS-Tween. After incubation for 15 min in 10 mM glycine, PBS-Tween, images were captured at 100x magnification using an excitation wavelength of 486 nm and 632 nm, and emission wavelengths of 501 nm and 647nm, respectively, with a DM-IRE2 confocal laser scanning microscope (Leica Microsystems Heidelberg GmbH, Germany). Negative controls consisted of biofilms being incubated with either no primary anti-rSslE antibody or no Alexa Fluor 633 secondary antibody.

### Immunoelectron microscopy

Overnight cultures were incubated for 30 min in 20 mM citrate phosphate buffer at either pH 4.0 or 5.0 and then washed in the same buffer before fixing with the 3% (w/v) paraformaldehyde for 1 hr. Cells were then loaded onto a Glow discharged carbon coated Ni grid (Agar Scientific) for 10 mins, washed with 50 mM glycine and then blocked with 1% (v/v) Natural Donkey Serum (Jackson Immunoresearch) for 30 min. This was then incubated for 1 hr with primary polyclonal rSslE antibody (rabbit; Invitrogen) diluted 1:100 in blocking buffer incubation, washed five times for 3 min each with 0.05% Natural Donkey Serum and then incubated for 1 hr with gold-conjugated secondary antibody (donkey; Jackson Immunoresearch) diluted 1:100 in blocking buffer. Washing was repeated with buffer only ddH_2_O. The sample was negatively stained with 2% (w/v) uranyl acetate for 30 s, followed by two quick washes with double-distilled water. The grid was air dried, and images were recorded on a JEM1230 (JEOL-Japan) at 80KV with a Morada CCD camera, iTEM software (EMSIS).

### Protein purification

Recombinant SslE (residues 67-1497; numbered based on mature sequence) was expressed and purified as described previously^12^. Likewise, SslE NT1 (residues 67-211), NT2 (residues 230-425), NT1-NT2 (residues 67-425), NT3-M60 (residues 426-1497) and RgpB-CTD were transformed into *E. coli* SHuffle cells (SslE; New England Biolabs) or BL21 (DE3) (RgpB-CTD; New England Biolabs), grown at 37°C in LB media (rSslE, RgpB-CTD: 100 μg/ml ampicillin; NT1, NT2, NT1-NT2, NT3-M60: 100 μg/ml kanamycin) and expression induced with 0.5 mM isopropyl-d-1-thiogalactopyranoside (IPTG) at an OD_600nm_ of 0.6, followed by growth overnight at 18°C. Cells were resuspended in 20 mM Tris–HCl pH 8, 200 mM NaCl, lysed by sonication and purified using nickel affinity chromatography followed by gel filtration with either a Superdex 75 (NT1, NT2, RgpB-CTD) or 200 (rSslE, NT1-NT2, NT3-M60) column (GE Healthcare).

### SEC-SAXS

SAXS data were collected on beamline B21^52^ at Diamond Light Source (DLS, Oxford, UK), United Kingdom at 25°C. 60 μl of rSslE (10 mg/ml) in 20 mM Tris–HCl pH 8, 200 mM NaCl was applied to a KW403-4F column (Shodex) at 0.16 ml/min, preequilibrated in 20 mM citrate phosphate buffer, 200 mM NaCl at either pH 4.4 or 7.4, and SAXS data were measured over a momentum transfer range of 0.003 < *q* < 0.44 Å^−1^. Peak integration and buffer subtraction were performed in CHROMIXS^53^. The radius of gyration (*R*_*g*_) and scattering at zero angle [*I(0)*] were calculated from the analysis of the Guinier region by AUTORG^53^. The distance distribution function [*P(r)*] was subsequently obtained using GNOM^53^ yielding the maximum particle dimension (*D*_max_). *Ab initio* low resolution shape restoration was carried out by calculating 20 models in DAMMIF^53^, which were subsequently averaged using DAMAVER^53^ and used as a staring model for refinement in DAMMIN^53^. An additional 20 models were also calculated and averaged in DAMAVER^53^. CRYSOL^53^ was used to compare final rSslE TEM envelopes and models against the solution SAXS curve. Processing and refinement statistics can be found in **Supplementary Table 2**.

### TEM single particle analysis

4 μl of rSslE (625 nM) in 50 mM Tris-HCl pH 8.0, 150 mM NaCl was applied to previously glow-discharged 300 mesh continuous carbon-coated copper grids (Agar Scientific Ltd) for 1 min and blotted for excess liquid. 4 μl of 2% (v/v) uranyl acetate was applied for staining for 1 min. The excess liquid was blotted and left to dry. Data was acquired using a JEOL JEM-2100 plus TEM operating at 200 kV equipped with a OneView 16 Megapixel camera (Gatan). 50 micrographs were collected at a nominal magnification of 50,000× with a pixel size of 2.1 Å/pixel and a range of defocus from 1 to 3 μm. Data were processed using Relion 3.1^54^. Defocus and astigmatism parameters were estimated using CTFFIND4^55^ in Relion 3.1. An initial dataset of 41,752 particles were autopicked using 2D class averages generated using approx. 1500 manually picked particles as reference. After a few rounds of 2D classification ignoring CTF until the first peak, 10,988 particles were taken forward for an initial model generation using 3D initial model in Relion 3.1^54^. Following initial model generation, a few rounds of 3D classification followed by 3D refinement was carried out. The final model was refined to 22 Å using the gold standard FSC (0.143 criterion). PHYRE2^56^ was used to analyse the SslE sequence (residues 67-1497; UniProt ID E0IW31) and generate a homology model for the C-terminal M60 domain (residues 962-1415) based on residues 421-894 of *P. aueriginosa* IMPa (PDB ID code 5kdv; 100% confidence, 21% identity)^19^. This was then docked into the rSslE TEM envelope using UCSF CHIMERA^57^.

### Solution NMR spectroscopy

NMR measurements for rSslE were performed at 25°C on a 100 μM ^2^H^15^N-labelled sample in 50 mM NaPO_4_ pH 7.4, 100 mM NaCl, 10 % D_2_O. NMR measurements for SslE NT1 (0.6 mM), SslE NT2 (1.3 mM) and SslE NT1-NT2 (0.8 mM) were performed at 25°C on ^15^N-labelled samples in 50 mM NaPO_4_ pH 7.0, 100 mM NaCl, 10 % D_2_O. NMR measurements for RgpB-CTD (0.3 mM) were performed at 37°C on a ^15^N-labelled sample in 20 mM NaPO_4_ pH 6.0, 100 mM NaCl, 10% D_2_O. Transverse relaxation optimized spectroscopy (TROSY) based ^1^H^15^N-HSQC experiment and *T*_1_ and *T*_2_ relaxation times for rSslE were recorded on a Bruker Avance III HD 950, equipped with a TXI cryoprobe. TROSY ^1^H^15^N-HSQC experiment for SslE NT1-NT2 was recorded on a on a Bruker Avance III HD 800, equipped with a TCI cryoprobe. Standard ^1^H^15^N-HSQC spectra of SslE NT1, SslE NT2 and RgpB-CTD were recorded on a Bruker Avance III HD 700, equipped with a TCI cryoprobe. Data were processed in NMRPIPE^58^ and analysed/visualized with ANALYSIS^59^ and NMRVIEW^60^.

### SAXS fibrillation analysis

SAXS data were collected on beamline B21^52^ at DLS (Oxford, UK) at 20°C. Immediately prior to data collection, 10 mg/ml rSslE in 20 mM Tris–HCl pH 8, 200 mM NaCl was buffer exchanged into 20 mM citrate-phosphate buffer, pH 4.4 using a PD10 column (GE Healthcare) and the flow through was used as a buffer reference. Using a peristaltic pump and while constantly stirring, 5 ml of rSslE (0.7 mg/ml) was circulated through the SAXS imaging cuvette and data were collected every 30 min over a momentum transfer range of 0.004 < *q* < 0.4 Å^−1^, with the initial scattering data captured at 30 min after initiating fibre growth. Data collected over the course of 11 hrs, consisting of 22 profiles, was decomposed using COSMiCS^30^, which utilizes MCR-ALS^61^ to perform model free decomposition of the entire SAXS data set. The SEC-SAXS curve collected at pH 4.4 was also included in the dataset as the representative state of the protein at time 0 s, *i.e.* before initiating fibrillation. The time 0 s curve was selected as one of the initial estimates and the selectivity restraint was used to ascertain that the curve had no contribution from the other species. In addition, non-negativity restraint was imposed for both the SAXS profiles and concentrations using FNNLS algorithm^62^. Before COSMiCS analysis, SAXS data were scaled according to the large angle data to enhance the decomposition capacity of the approach. Although COSMiCS was run assuming a two-component system as suggested by Principal Component Analysis (PCA), a three-species run was also performed. The convergence criterion of < 0.01 % change in lack of fit was used with 1,000 maximum allowed iterations. The analysis of the resulting COSMiCS curves was performed with the ATSAS suit of programs^53^. Fractal analysis was carried out using SASVIEW (http://www.sasview.org/).

### SAXS fibre modelling

The programs ATSAS^53^ and Scatter were used to obtain a cross-sectional radius of gyration *Rc* from the 10 hr post-fibre induction scattering profile and from this the cross-sectional *P*(*r*)_*C*_ was calculated. BODIES^53^ was then used to approximate the geometric shape of rSslE fibres, which suggested a cylindrical shape with radius of 3.93 nm and a height of 16.5 nm (χ^2^ 1.6). These dimensions were then used in DAMMIN^63^ with data from the q range 0.03 to 0.20 to create *ab initio* shapes of the SslE fibril repeat. 20 individual jobs were run, generating 20 independent models and were subsequently averaged using DAMAVER^53^ and then refined 5 times in DAMMIN^63^. A bead model was produced from the rSslE negative-stain TEM envelope using the program EM2DAM^53^, the NT1 bead region was removed and then this was superposed onto the DAMMIN fibre bead model using SUPCOMB^53^. Processing and refinement statistics can be found in **Supplementary Table 2**.

### TEM fibre analysis

rSslE at 1 mg/ml was incubated in 100 mM citrate phosphate pH 3.8 overnight at room temperature while shaking at 180 rpm. This was centrifuged at 15,000 g for 10 min, the top 80% of buffer discarded and then 4 μl of the remaining sample was applied to a previously glow-discharged 300 mesh continuous carbon-coated copper grids (Agar Scientific Ltd) and immediately blotted for excess liquid. 4 μl of 2% (v/v) uranyl acetate was applied for staining for 10 seconds. The excess liquid was blotted and left to dry. Data was acquired using a JEOL JEM-1230 TEM operating at 80 kV equipped with a Morada 2k CCD camera system and its iTEM software package (Olympus Europa, UK). Micrographs were collected at a nominal magnification of 80,000x with a pixel size of 5.96 Å/pixel.

### Solid state NMR spectroscopy

rSslE at 20 mg/ml was buffer exchanged into 10 mM citrate phosphate pH 4.0 using a PD10 column (Sigma) and then incubated overnight at room temperature while shaking at 180 rpm. This was centrifuged at 15,000 g for 10 min, the top 80% of buffer was discarded and then the remaining sample was used for solid state NMR analysis. Experiments were performed using a Bruker Neo Console operating at 850 MHz ^1^H frequency with a 3.2 mm E-Free probe in HC mode spinning at a rate w_r_= 15 kHz. A standard CP excitation ^13^C-^13^C DARR experiment was acquired. The direct dimension was acquired for 16.4 ms with ~55 kHz SPINAL-64 decoupling. 256 rows with 256 coadded transients and a recovery delay of 2.5 s were acquired using Time-Proportional Phase Increment (TPPI) with a 22.2 μs dwell (45 kHz sweep width, 2.84 ms total evolution) for a total of 45.5 hrs total acquisition time. The applied power was adjusted so that ^1^H and ^13^C hard pulses were both 4 μs (w_1(H,C)_= 62.5 kHz). The initial carbon excitation was achieved with 1.5 ms of ramped Hartmann-Hahn CP, where the optimal polarization transfer was found at w_1C_= ~70 kHz and w_1H_= ~55 kHz with an upwards linear ramp from 70-100% on the ^1^H channel. Homonuclear Carbon mixing was achieved with 50 ms of DARR mixing (w_1H_= w_r_= 15 kHz).

## Acknowledgements

PC was supported by the Academy of Medical Sciences and Wellcome Trust (SBF002/1150) and SR was supported by the Medical Research Council (MR/R017662/1), awarded to JAG. SW and BD were supported by China Scholarship Council and Biotechnology and Biological Sciences Research Council studentships, respectively, awarded to JAG and MAC. LS was supported by the Leverhulme Trust (RPG-2017-222), awarded to JAG. KF was supported by a PhD studentship from Queen Mary University of London and VCD was supported by a start-up grant from Queen Mary University of London. This work was also supported by the Labex EpiGenMed, an « Investissements d’avenir » program (ANR-10-LABX-12-01) awarded to PB. The CBS is a member of France-BioImaging (FBI) and the French Infrastructure for Integrated Structural Biology (FRISBI), 2 national infrastructures supported by the French National Research Agency (ANR-10-INBS-04-01 and ANR-10-INBS-05, respectively). We thank the beamline scientists at B21 of the Diamond Light Source, United Kingdom. This work was also supported by the Francis Crick Institute through provision of access to the MRC Biomedical NMR centre. The Francis Crick Institute receives its core funding from Cancer Research United Kingdom (FC001029), the United Kingdom Medical Research Council (FC001029), and the Wellcome Trust (FC001029). We also thank the Centre for Biomolecular Spectroscopy at King’s College London for additional NMR access, funded by the Wellcome Trust and British Heart Foundation (ref. 202767/Z/16/Z and IG/16/2/32273 respectively). The UK 850 MHz solid-state NMR Facility used in this research was funded by EPSRC and BBSRC (contract reference EP/T015063/1), as well as the University of Warwick including via part funding through Birmingham Science City Advanced Materials Projects 1 and 2 supported by Advantage West Midlands (AWM) and the European Regional Development Fund (ERDF). Cellulose discs were gifted by Prof. Tom Ellis and Kenneth T. Walker (Imperial College London).

## Author Contributions

Conceived and designed the experiments: PC, SW, SR, KF, SS, CE-G, AS, GM, BD, LMS, SL, DI, PB, VCD, JAG. Performed the experiments: PC, SW, SR, KF, AS, SS, LC, CE-G, GM, BD, LMS, SL, DI, JJ, GHC, VCD, JAG. Analyzed the data: PC, SW, SR, KF, AS, SS, CE-G, GM, BD, SL, DI, TF, PB, VCD, JAG. Contributed reagents/materials/analysis tools: SS, SL, PB, VCD, JAG. Wrote the paper: PC, SW, SR, KF, AS, SS, LC, CE-G, GM, BD, LMS, SL, DI, TF, JJ, GHC, MAC, PB, VCD, JAG.

