## Supplementary material for "Molecular and cellular insight into *Escherichia coli* SslE and its role during biofilm maturation": Suplementary Information

### Supplementary Methods

##### SslE secretion

*E. coli* W or H10407 were grown on LB agar with 50 µg/ml kanamycin for *Δssle* mutants and 50 µg/ml kanamycin, 25 µg/ml chloramphenicol for *Δssle::pCPC1* (*sslE*) and *Δssle::pCPC2* (*sslEΔM60*). Single colonies were resuspended in 10 ml LB (with appropriate antibiotics) and incubated overnight at 37°C with shaking (200 rpm). Cells were centrifuged to separate out the media (supernatant) and pellet (whole cells) and 3 µl of samples were resuspended in 17 µl of 1x NuPAGE LDS Sample Buffer (ThermoFisher), 5% (v/v) β-Mercaptoethanol. Samples were run on a Criterion 4-20% SDS-PAGE gel (Bio-Rad), followed by transfer onto a PVDF membrane using the semi-dry Invitrogen Power-Blotter and Power Blotter transfer blotting solution. The membrane was blocked with 1% (w/v) BSA, PBS-Tween for 1 hr at room temperature, and then incubated overnight at 4°C with polyclonal anti-rSslE antibody (rabbit; Invitrogen) or monoclonal anti-DsbA (mouse; Invitrogen), diluted 1:1000 using 0.5% (w/v) BSA, PBS-Tween incubation buffer. After three, 5 min washes with incubation buffer, membranes were incubated for 1 hr at 37°C with either anti-rabbit or anti mouse secondary antibody conjugated to HRP (1:2000 dilution; Invitrogen) for 1 hr at 25°C and then treated with enhanced chemiluminescence substrate (ECL; Pierce) before detection.

**Macrocolony biofilm growth**

5 µl of wild-type *E. coli* W or H10407 strains and their derivatives were grown on LB agar containing either 0.8 mg/ml Congo red dye or 50 µM calcofluor white at 37°C for either 24 or 96 hrs. 50 µg/ml kanamycin was included for *Δssle* mutants and 50 µg/ml kanamycin, 25 µg/ml chloramphenicol was included for *Δssle::pCPC1* (*sslE*) and *Δssle::pCPC2* (*sslEΔM60*). Colonies were visualised with white light and either epi-red (Congo red) or UV (calcofluor white) light.

**Assessment of biofilm growth**

FilmTracer FM1-43 dye (Life Technologies) was added to the BioFlux 200 input reservoir mixed with prewarmed LB to a final concentration of 0.3 µM. The heated stage holding the BioFlux plate was then moved to a DM-IRE2 confocal laser scanning microscope (Leica Microsystems Heidelberg GmbH, Germany) for image acquisition with the Leica Microsystems Confocal Software (version 2.61 Build 1537). The fluorophore was excited at 472 nm and emission collected at 580 nm as recommended by the manufacturer.

**Bacterial culture ring assay**

10 ml of LB containing 0.8 mg/ml congo red dye, adjusted to either pH 5.0 or 7.0 with HCl, were inoculated with wild-type *E. coli* W and derivatives (including 50 µg/ml kanamycin for *Δssle* mutant and 50 µg/ml kanamycin, 25 µg/ml chloramphenicol for *Δssle::pCPC1* (*sslE*) and *Δssle::pCPC2* (*sslEΔM60*)). These were incubated overnight at 37°C while shaking (200 rpm) and the presence of ring formation on the glass tube was visually assessed.

**Recombinant protein ring assay**

20 µl of rSslE, NT1-NT2 or NT3-M60 (62.5 µM) in 10 mM Tris-HCl pH 8, 100 mM NaCl buffer were diluted to 2 ml in 100 mM citrate phosphate buffer at pH 4.0, 4.2, 4.4, 4.6, 4.8, 5.0, 5.4, 5.8, 6.2, 6.6 and 7.0. These were transferred to glass tubes, incubated overnight at 37°C while shaking (200 rpm) and the presence of ring formation on the glass tube was visually assessed. Buffer from tubes containing rings formed at pH 4.0 was then removed without disturbing the rings and 3 ml of 100 mM citrate phosphate buffer over the same pH range was added. These were left overnight with shaking at 37°C and solubilization of the rings were visually assessed the next day.

**ThT binding**

15 µl of 50 µM rSslE in 10 mM Tris-HCl pH 8, 100 mM NaCl buffer was mixed with 1.5 µl of 1 mM Thioflavin T (ThT) dye and transferred to a 96-well plate. 100 mM citrate phosphate buffer between pH 4.0 and 8.0 was added to a final volume of 150 µl and fluorescence data was collected (excitation/emission 438/480 nm) at 37°C with shaking every 15 min over 24 hrs using a BMG CLARIOstar plate reader.

**ATR FT-MIR spectroscopy**

FT-MIR spectroscopy measurements were acquired using a Perkin Elmer Frontier spectrometer (Perkin Elmer, Buckinghamshire, UK) in attenuated total reflectance (ATR) mode, fitted with a Golden Gate ATR accessory (Specac Ltd., Kent, UK) with a diamond crystal. Samples were enclosed in a sealed Plexiglas container within a nitrogen filled atmosphere at 23 ± 1 °C with additional calcium chloride dihydrate pellets (Sigma). Spectra were acquired using an infrared light source and a DTGS (deuterated triglycine sulphate) detector over a spectral range of 500 to 4000 $cm^{-1}$ with a 4 $cm^{-1}$ spectral resolution (filling factor=2) and 64 co-added scans. 5 µl rSslE solution, as either soluble monomer in PBS buffer (pH 7.4) or fibres in 100 mM phosphate-citrate buffer (pH 4.4), was deposited and dried onto the ATR crystal (n=3 per protein solution). Spectral data were background subtracted and baseline corrected using the Spectrum One software package (version 6, Perkin Elmer, Buckinghamshire, UK). To deconvolve the spectra for curve fitting second derivative spectra were calculated, using the Savitzsky-Golay method with a 9-point window and a third-degree polynomial, to identify the positions of overlapping component bands. Voigt peak shapes combined with a linear background were fitted to the amide I region (1580-1710 $cm^{-1}$) of the fibre sample FT-MIR spectrum, fixing the peak positions obtained via second derivative spectra whilst leaving other parameters free to adjust iteratively. Integrated deconvolved band intensities correspond to the relative proportions of each absorption band and secondary structure. Curve fitting was performed using the Lmfit package (version 1.0.0, Zenodo). To assign secondary structures to band components, frequencies in the range 1580-1610 $cm^{-1}$ were assigned to side chains and aggregated β strands, 1611–1630 and 1697-1703 $cm^{-1}$ were assigned to intermolecular β-sheet structures, 1630–1637 $cm^{-1}$ to intramolecular β-sheet structures, 1638-1655 $cm^{-1}$ to random coil structures, 1656–1662 $cm^{-1}$ to α-helical structures and 1662–1687 $cm^{-1}$ to β-turns[1, 2, 3, 4].

**Far-UV CD spectroscopy**

Far-UV CD spectra were measured on a Chirascan (Applied Photophysics) spectropolarimeter thermostated at 25°C. rSslE (10 mg/ml) in 20 mM Tris–HCl pH 8, 50 mM NaCl was diluted 200-fold to 0.05 mg/ml in 10 mM MOPS at either pH 4.4 or 7.4. Spectra were recorded from 260 to 190 nm, at 0.2 nm intervals, 1 nm bandwidth, and a scan speed of 50 nm/min. Three accumulations were averaged for each spectrum. Deconvolution of data was performed using the BESTSEL server[5].

**Mass spectrometry**

Ring aggregates formed from either bacterial cultures or recombinant SslE were scraped from glass tubes and resuspended in 0.5 ml 100 mM citrate phosphate buffer, pH 4.0. Samples were resolved using SDS PAGE and after staining, SslE fibre bands retained in the wells were excised and incubated with 10 mM dithiothreitol at 56°C and then alkylated with 55 mM iodoacetamide at room temperature. Sample were digested using 1:20 dilution of bovine trypsin incubated in a shaking heat block at 37°C for 16 hrs. Peptides were extracted with aqueous dehydration/hydration using acetonitrile and 50 mM triethylammonium bicarbonate, pooled and dried. Samples were resuspended in 2% (v/v) acetonitrile, 0.05% (v/v) formic acid and peptides were resolved by reversed phase chromatography on a 75 µm C18 Pepmap column (50 cm length) using a three-step linear gradient of 80% acetonitrile in 0.1% formic acid (U3000 UHPLC NanoLC system; ThermoFisherScientific, UK). The gradient was delivered to elute the peptides at a flow rate of 250 nl/min over 60 min starting at 5% B (0-5 min) and increasing solvent to 40% B (5-40 min) prior to a wash step at 99% B (40-45 min) followed by an equilibration step at 5% B (45-60 min). The eluate was ionised by electrospray ionisation using an Orbitrap Fusion Lumos (ThermoFisherScientific, UK) operating under Xcalibur v4.1.5. The instrument was first programmed to acquire using an Orbitrap-Ion Trap method by defining a 3 s cycle time between a full MS scan and MS/MS fragmentation. Orbitrap spectra (FTMS1) were collected at a resolution of 120,000 over a scan range of m/z 375-1500 with an automatic gain control (AGC) setting of 4.0e5 with a maximum injection time of 35 ms. Monoisotopic precursor ions were filtered using charge state (+2 to +7) with an intensity threshold set between 5.0e3 to 1.0e20 and a dynamic exclusion window of 35 secs ± 10 ppm. MS2 precursor ions were isolated in the quadrupole set to a mass width filter of 1.6 m/z. Ion trap fragmentation spectra (ITMS2) were collected with an AGC target setting of 1.0e4 with a maximum injection time of 35 ms with CID collision energy set at 35%. Data were processed using Proteome Discoverer (v2.2; ThermoFisher) to search against Uniprot Swissprot All Taxonomy (561,911 entries) and the sequence of SslE (Uniprot Accession number - E3PJ90) with Mascot search algorithm (v2.6.0; www.matrixscience.com) and the Sequest search algorithm[6]. Precursor mass tolerance was set to 20 ppm with fragment mass tolerance set to 0.8 Da with a maximum of two missed cleavages. Variable modifications included: Carbamidomethylation (Cys) and Oxidation (Met). Database generated files (.msf) were uploaded in to Scaffold software (v 4.10.0; www.proteomesoftware.com) for visualisation of fragmentation spectra.

**SslE fibre immunoblot**

Ring aggregates formed from rSslE were scraped from glass tubes and resuspended in 0.5 ml 100 mM citrate phosphate buffer, pH 4.0, centrifuged at 15,000 g and then the top 950 µl solution was carefully removed and discarded. This was followed by three rounds of addition of 950 µl 100 mM citrate phosphate buffer at pH 4.0, centrifugation at 15,000 g and the top 950 µl discarded. The final 50 µl sample was mixed with 1x NuPAGE LDS Sample Buffer (ThermoFisher), 5% (v/v) β-Mercaptoethanol and incubated at 100°C for 5 min prior loading. This was run on a Criterion 4-20% SDS-PAGE gel (Bio-Rad), followed by transfer onto a PVDF membrane using the semi-dry Invitrogen Power-Blotter and Power Blotter transfer blotting solution. The membrane was blocked in 1% (w/v/) BSA, PBS-Tween for 1 hr at room temperature followed by the addition of 1:2000 dilution mouse anti-$His_6$ antibody (Sigma) in 0.5% (w/v/) BSA, PBS-Tween incubation buffer for 2 hrs. After 5 rounds of washing with incubation buffer, the membrane was incubated with 1:2000 anti-mouse HRP-conjugated antibody (Sigma) for 1hr, followed by 5 further washes and then treatment with ELC substrate (Peirce) before detection.

**RT-MALS**

1 ml of rSslE (1 mg/ml) in 25 mM citrate phosphate buffer pH 4.0 was incubated overnight at room temperature while shaking at 180 rpm. The sample was then centrifuged at 15,000 g and the top 850 µl of buffer was discarded. MALS experiments were then performed on beamline B21 at the DLS (Oxford, UK). 20 uL of SslE fibres were directly injected into the RT-MALS multiple times at a flow rate of 0.05 ml/min. Detectors were standardised through a direct injection using BSA. Data was analysed using ASTRA version 6.1.7. (Wyatt).

**Cellulose binding assay**

An Immulon 2-HB 96-well plate (VWR) was blocked for 1 hr at 25°C with 300 µl of 0.1 % (w/v) BSA in PBS-Tween and then washed once with 300 µl of incubation buffer (100 mM citrate-phosphate buffer pH 6.0, 0.05 % (w/v) BSA, 0.05 % Tween-20). One cellulose disc[7] was added to each well and then 200 µl of incubation buffer was added to cover the discs, followed by incubation for 5 min. The discs were washed twice in incubation buffer and then 200 µl of either 100 µM rSslE monomer, rSslE fibre (produced as described for immunoblotting) or RgpB-CTD control, all in incubation buffer, were added to the cellulose discs. The plate was incubated for 3 hrs at 25°C and then the discs were transferred carefully to new pre-blocked wells and washed with 200 µl incubation buffer. After repeating three times, the discs were incubated with anti-$His_6$ HRP conjugated antibody (ThermoFisher) diluted 1:2000 with incubation buffer for 1 hr at 24°C. The discs were removed again, and the washing step was repeated 3 times. Finally, the solution was removed from the wells and the discs were incubated with 150 µl of *o*-Phenylenediamine dihydrochloride (Sigma) for 30 min. The solution in the wells was stirred thoroughly, the discs were removed and then data were recorded at 450 nm.

#### Supplementary Figures

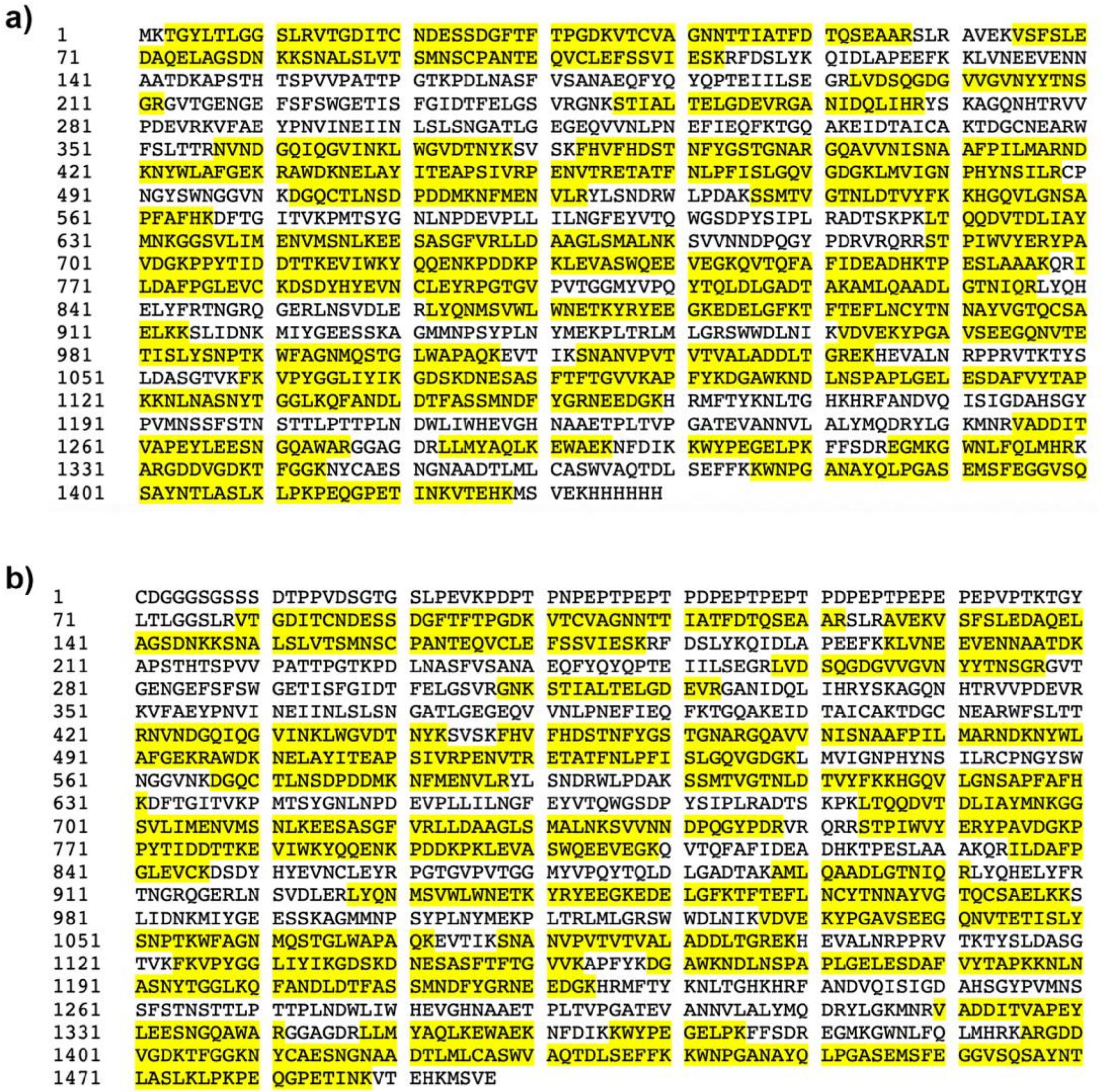


**Supplementary Fig. 1 | Mass Spec analysis of purified SslE aggregates. a,** Summary of peptides identified (yellow) in ring aggregates formed from recombinant rSslE at pH 4.4, with the rSslE sequence shown. **b,** Summary of peptides identified (yellow) in ring aggregates formed from *E. coli* W strain cultures at pH 5.0 with the sequence for mature SslE from *E. coli* W strain shown.

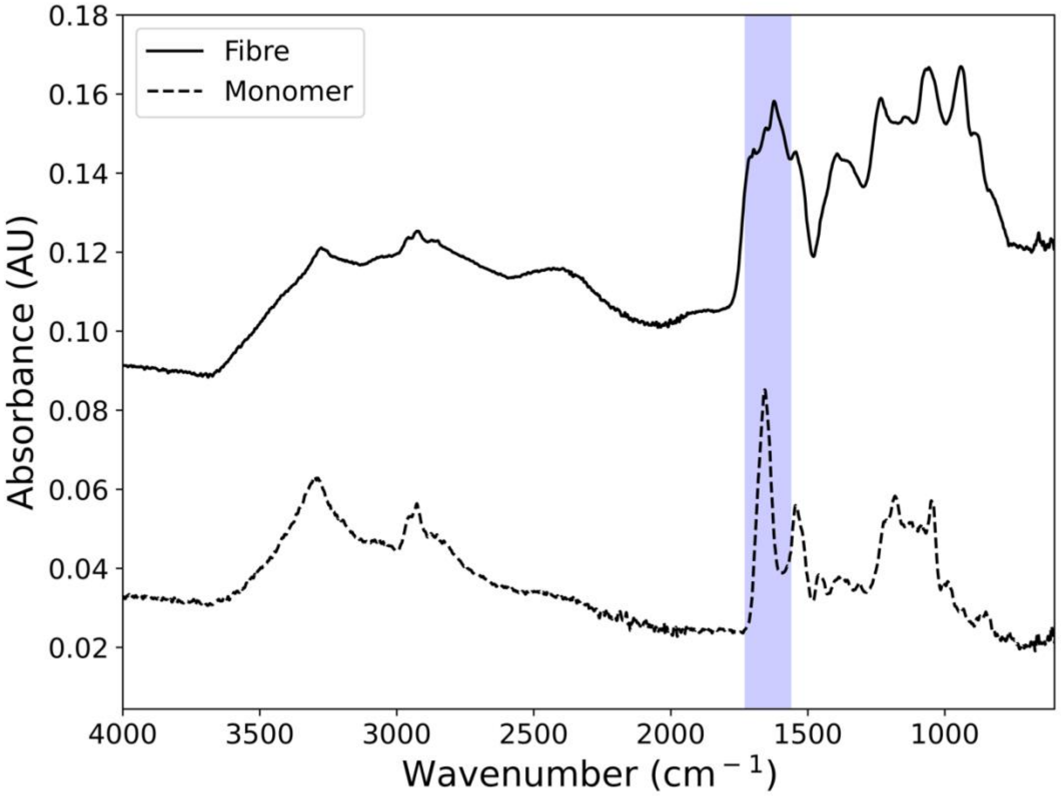


**Supplementary Fig. 2 | ATR FT-MIR spectra of rSslE in its monomeric and amyloid-like state.** Exemplar FT-MIR spectra for fibre (solid black line) and monomer (broken black line) specimens. The peak of the amide I absorption band (shaded blue region) is located at approximately 1624 and 1656 cm$^{-1}$ for the fibre and monomer specimens, respectively. The lower frequency amide I band for the fibre sample indicates an increase in anti-parallel β-sheet structure. Spectra are offset in the y-axis for clarity.

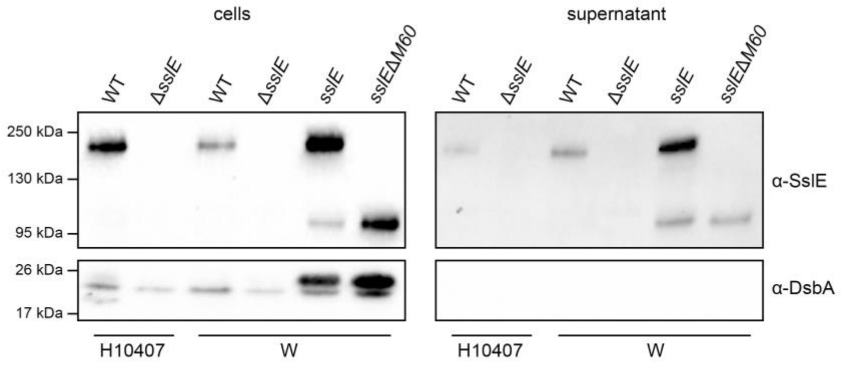


**Supplementary Fig. 3 | Secretion of SslE by *E. coli* W and H10407 and derivatives.** Immunoblot of overnight culture whole cell samples and culture supernatants detected with either anti-rSslE antibody or anti-DsbA antibody, which was used to confirm cell lysis had not occurred in supernatant samples. WT = wild-type; Δ*sslE* = *sslE* mutant; *sslE* = Δ*sslE::sslE*; *sslEΔM60* = Δ*sslE::sslEΔM60*.

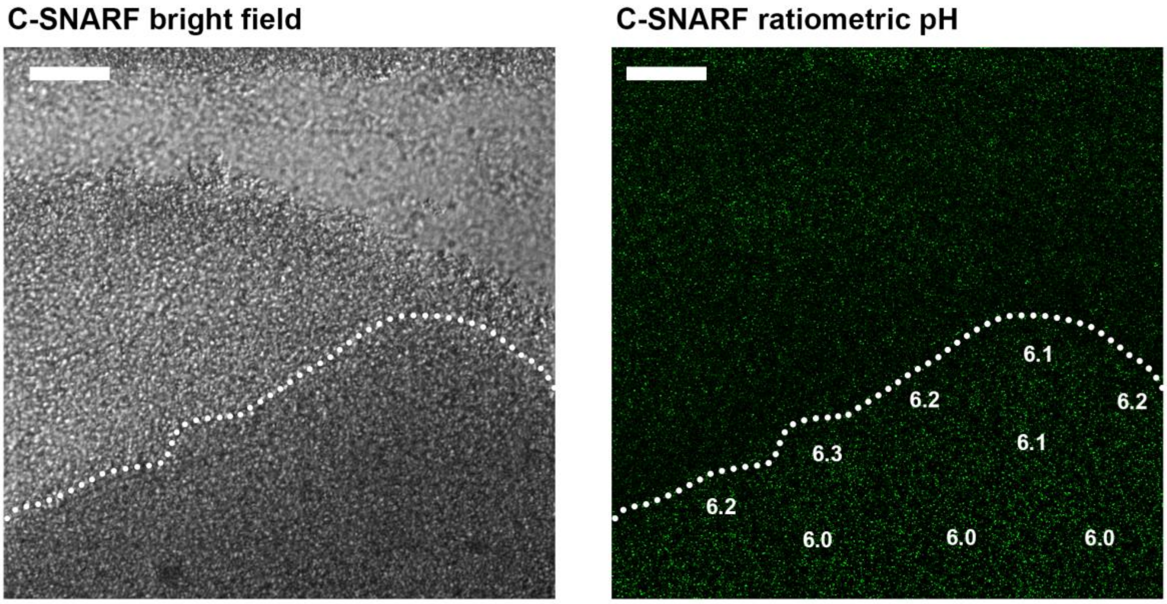


**Supplementary Fig. 4 | Analysis of pH across 24 hr old *E. coli* W biofilms.** The pH across biofilms grown for 24 hrs was monitored ratiometrically using C-SNARF-4 (green). pH values were calculated over 30 $\mu m^2$ boxes and pH values for representative regions are annotated. Dotted line outlines boundary of the biofilm mass. Scale bar represents 20 μm. All data are representative of at least three independent experiments.

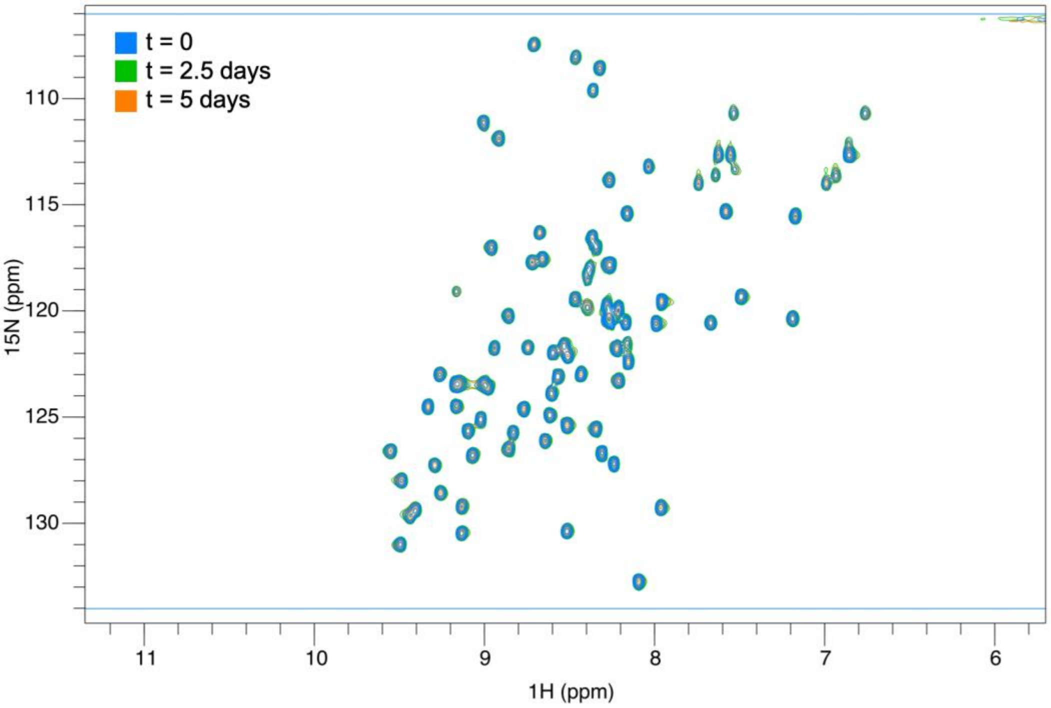


**Supplementary Fig. 5 | 2D $^{1}H^{15}N$-HSQC analysis of RgpB-CTD stability at pH 6.0.** Spectra were recorded after 0, 2.5 and 5 days incubation at 37°C. Excellent dispersion and narrow line widths can be observed for the majority of peaks suggesting this is a well folded domain. No new peaks are observed following 5 days incubation showing that the protein is stable in these conditions.

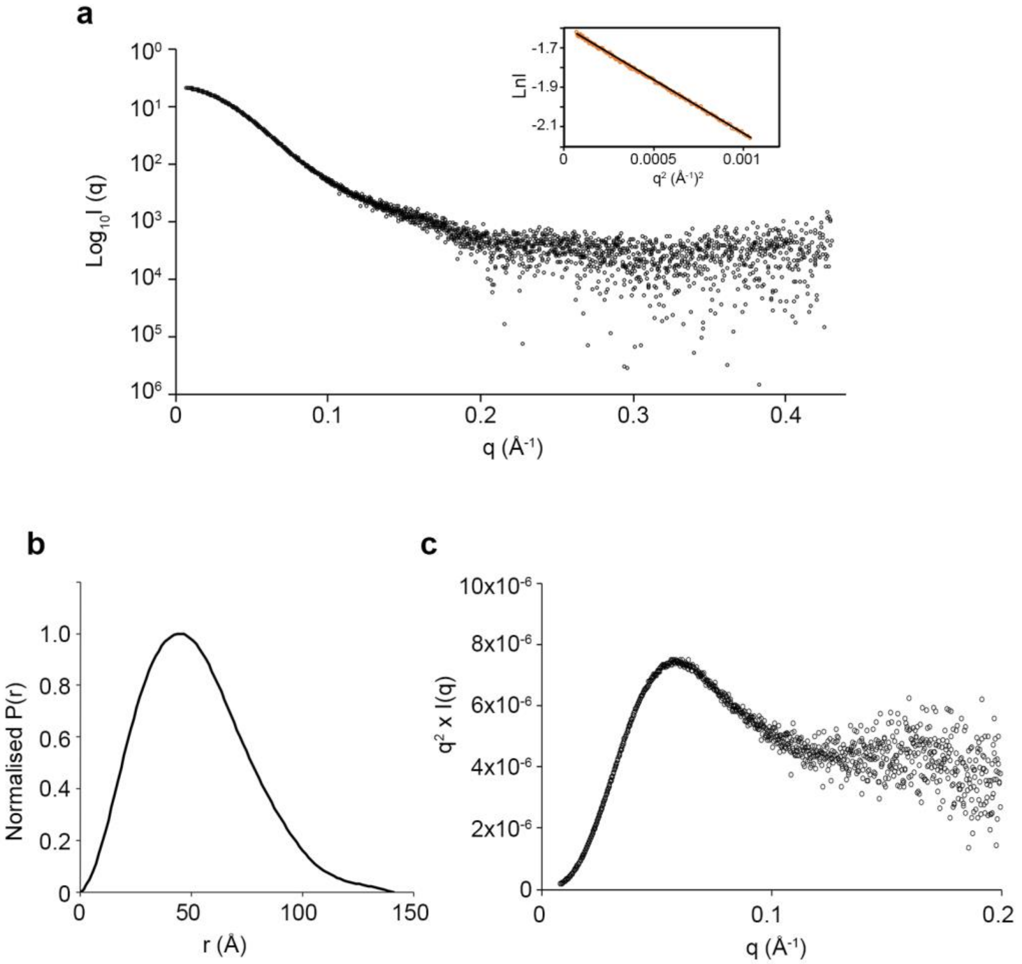


**Supplementary Fig. 6 | SAXS analysis of rSslE at pH 7.4. a,** Experimental scattering curve of rSslE (black open circles). Inset: Guinier Region (orange open circles) and linear regression (black line) for Rg evaluation. **b,** Shape distribution [P(r)] function derived from SAXS analysis for rSslE. **c,** Kratky plot indicates that rSslE has dynamic features in solution.

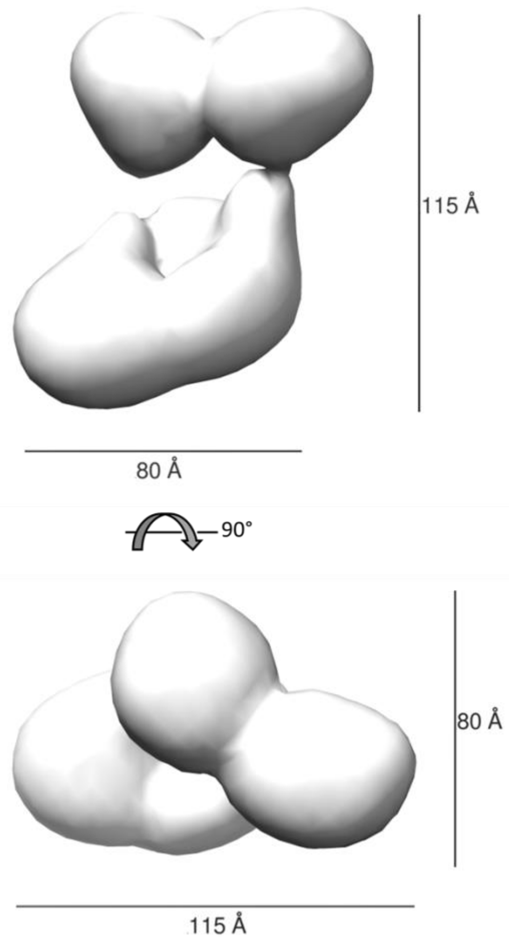


**Supplementary Fig. 7 | Overall dimensions of rSslE.** Negative-stain TEM map of rSslE is shown in two orientations with overall dimension of ~8×8×11.5 nm annotated.

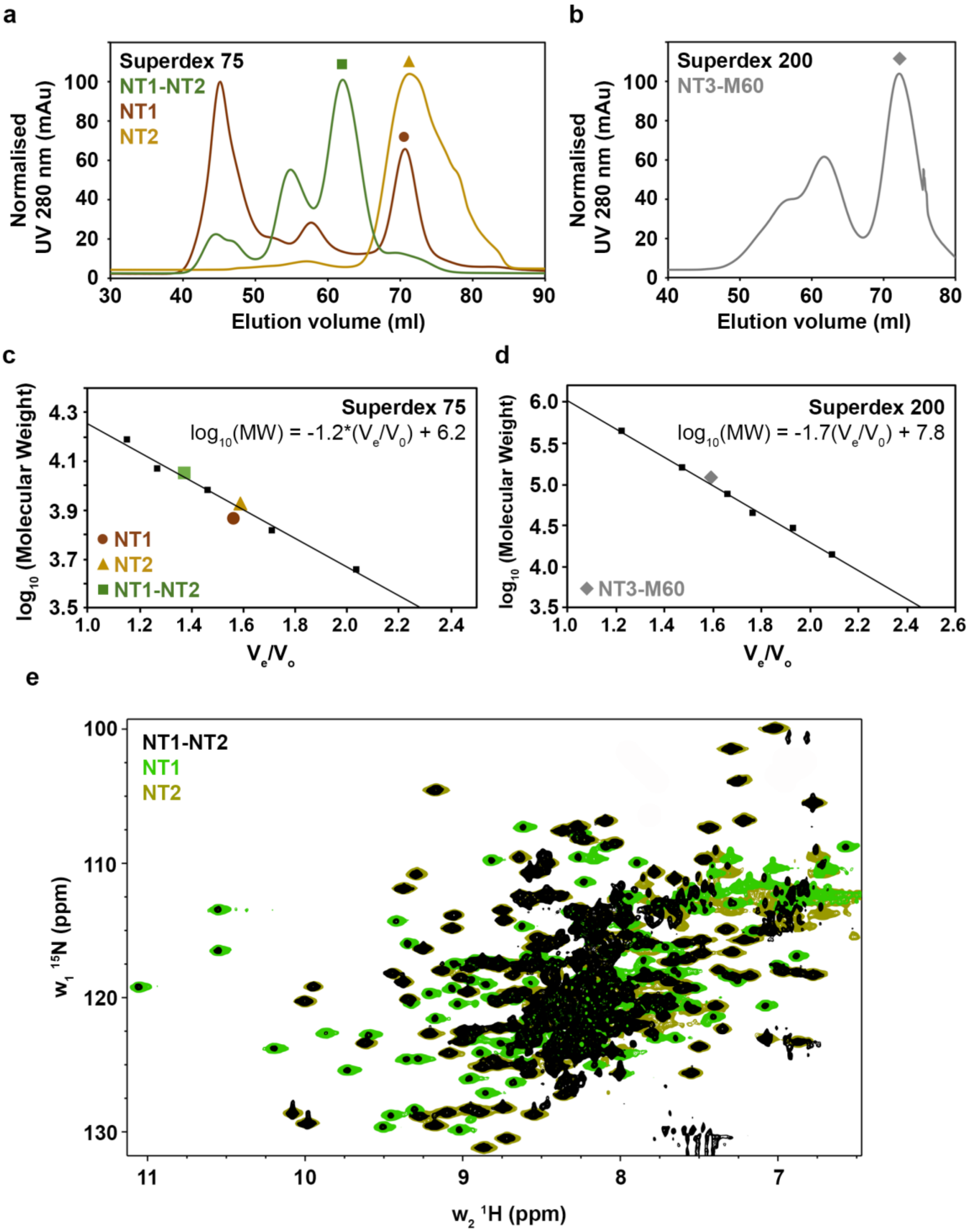


**Supplementary Fig. 8 | Folding analysis of SslE subdomain constructs. a-d,** Analytical size exclusion chromatography (SEC) of SslE NT1, NT2, NT1-NT2 and NT3-M60 constructs. **a,** Normalised chromatograms of NT1 (brown), NT2 (orange) and NT1-NT2 (green) injected onto a Superdex 75 column. **b,** Normalised chromatogram of NT3-M60 (grey) injected onto a Superdex 200 column. **c,** The $V_e/V_o$ (elution volume/column void volume) for the major peaks of NT1 (brown circle),

NT2 (orange triangle) and NT1-NT2 (green square) were plotted against their $Log_{10}$ molecular weights on a standard curve created using molecular weight standards (GE Healthcare). These profiles show that all constructs are likely monomeric in solution with calculated molecular weights of 21.8 kDa for NT1 (theoretic mass: 16.9 kDa), 20.7 kDa for NT2 (theoretic mass: 22.8 kDa) and 36.8 kDa for NT1-NT2 (theoretic mass: 40.1 kDa). **d,** The $V_e/V_o$ (elution volume/column void volume) for the major peak of NT3-M60 (grey diamond), NT2 (orange triangle) and NT1-NT2 (green square) was plotted against its $Log_{10}$ molecular weight on a standard curve created using molecular weight standards (GE Healthcare). This suggest that the construct is again a monomer in solution with a calculated molecular weight of 98.9 kDa (theoretic mass: 121.4 kDa). **e,** TROSY $^{1}H$-$^{15}N$ HSQC spectrum of $^{15}N$-labelled SslE NT1-NT2 (black) overlayed with $^{1}H$-$^{15}N$ HSQC spectra from the isolated $^{15}N$-labelled SslE NT1 (light green) and $^{15}N$-labelled NT2 (olive green). All spectra show well dispersed peaks, indicative of folded proteins, with good overlap between peaks from NT1 and NT2 with the NT1-NT2 spectrum.

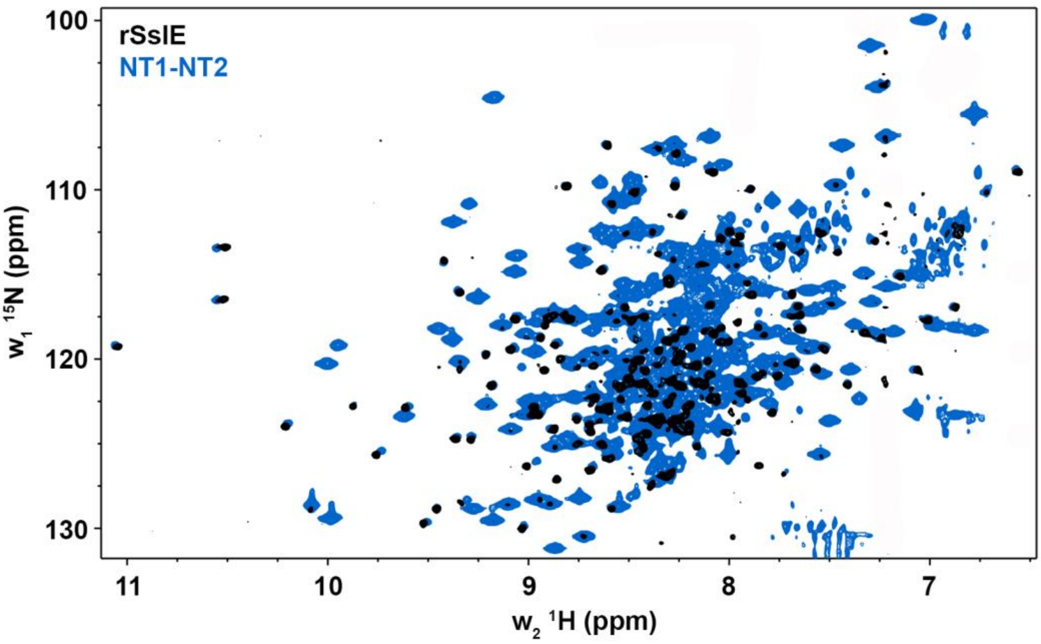


**Supplementary Fig. 9 | Comparison of 2D $^{1}H^{15}N$-HSQC spectra of rSslE and SslE NT1-NT2.** TROSY $^{1}H$-$^{15}N$ HSQC spectrum of $^{15}N$-labelled SslE NT1-NT2 (blue) overlayed with a TROSY $^{1}H$-$^{15}N$ HSQC spectrum from $^{2}H^{15}N$-labelled rSslE (black).

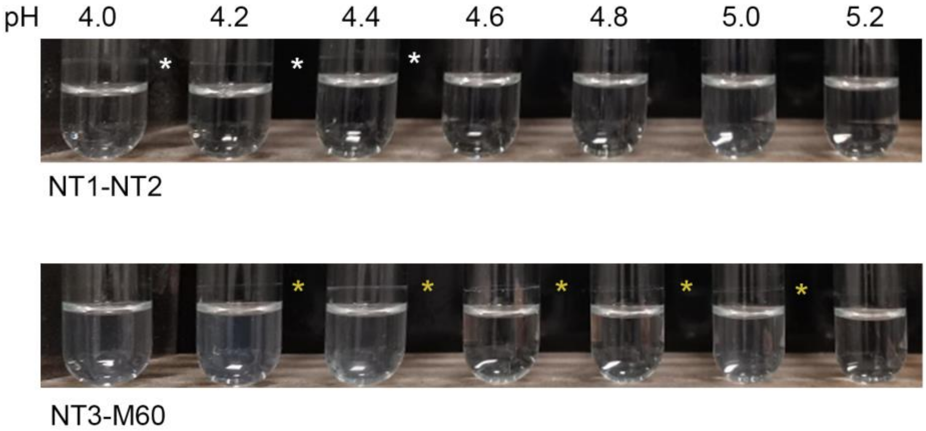


**Supplementary Fig. 10 | Recombinant NT1-NT2 and NT3-M60 aggregation assay.** Purified NT1-NT2 and NT3-M60 regions form rings of aggregated protein when incubated in citrate-phosphate buffer between pH 4.0 and 4.4 (NT1-NT2; white asterisks) and 4.2 and 5.0 (NT3-M60; yellow asterisks). The ring appears above the solution line due to shaking of the sample.

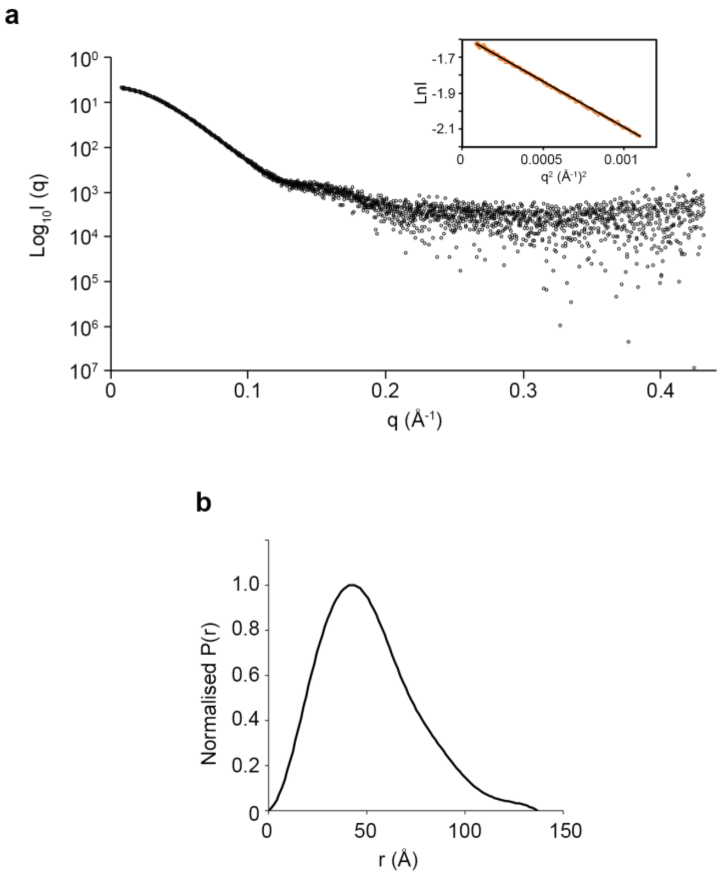


**Supplementary Fig. 11 | SAXS analysis of rSslE at pH 4.4. a,** Experimental scattering curve of rSslE (black open circles). Inset: Guinier Region (orange open circles) and linear regression (black line) for Rg evaluation. **b,** Shape distribution [*P(r)*] function derived from SAXS analysis for rSslE.

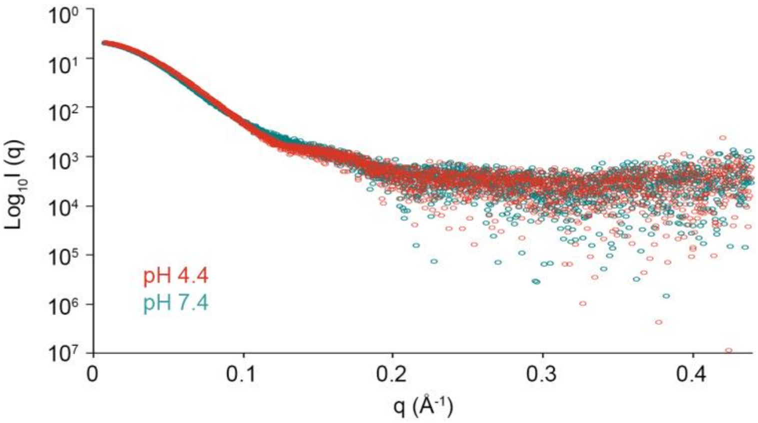


**Supplementary Fig. 12 | Comparison of rSslE SAXS scattering curves at different pH values.** SslE at pH 4.4 is red and at 7.4 is teal. Small differences are seen at q-values of approx. 0.13 Å$^{-1}$.

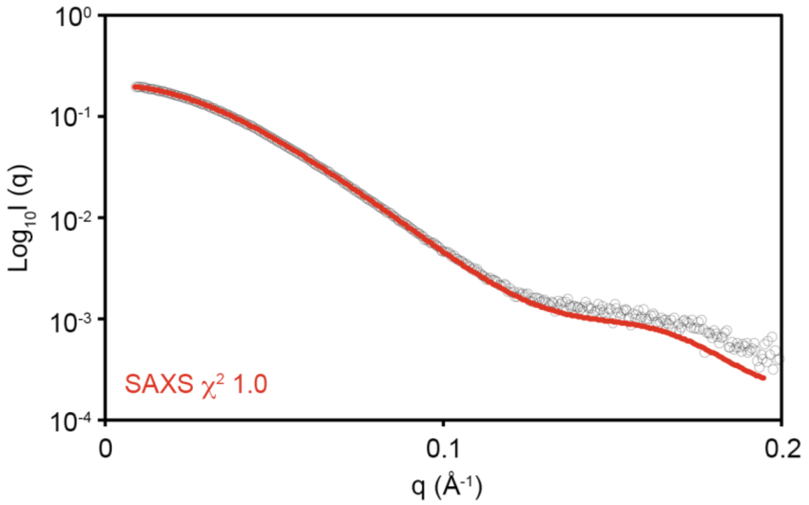


**Supplementary Fig. 13 | Fit of the rSslE SAXS bead model at pH 4.4 to SAXS data.** Bead model is shown as red line and SAXS data as black open circles, with $\chi^2$ of 1.0.

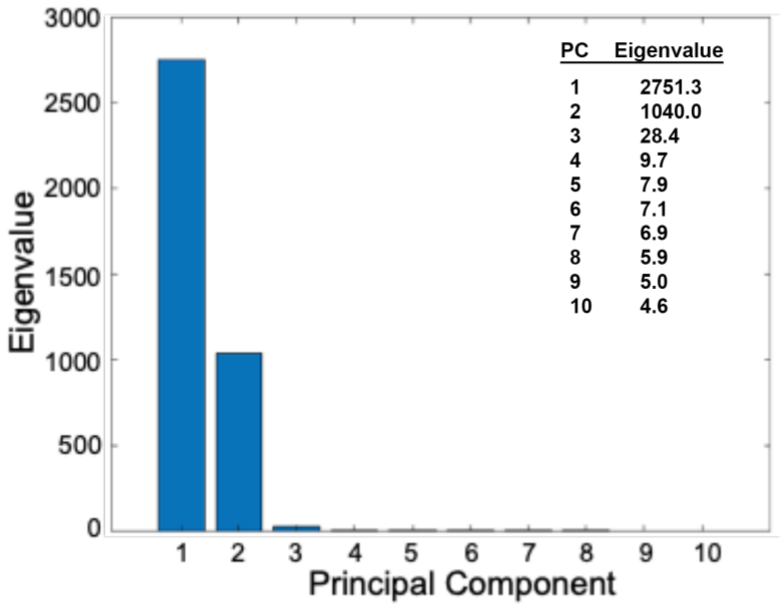


**Supplementary Fig. 14 | PCA analysis of rSslE SAXS fibrillation.** The Eigenvalues of the first ten principal components obtained after PCA analysis of the fibrillation SAXS dataset.

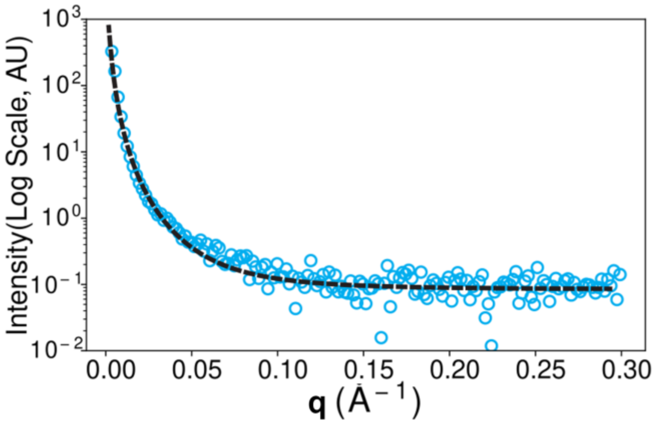


**Supplementary Fig. 15 | Fractal fit against rSslE SAXS fibrillation data.** The fractal-fit plot (black dashed line) fitted against the decomposed SAXS scattering profile from COSMiCS (black circles) using a mass-fractal model.

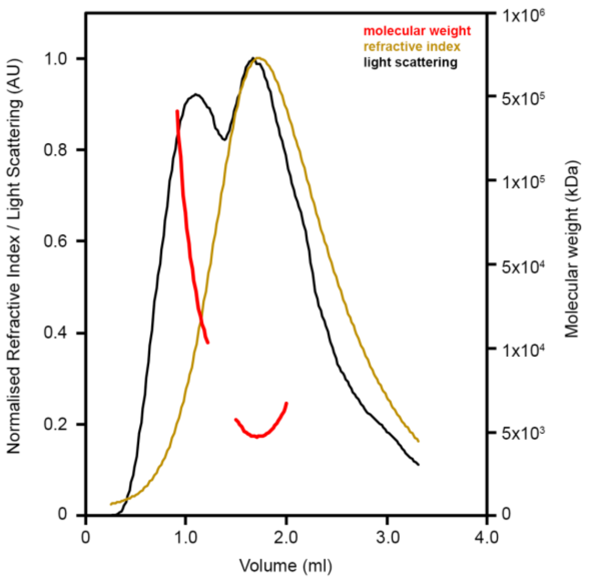


**Supplementary Fig. 16 | Mass analysis of rSslE aggregates using RT-MALS.** Two species are present, the lager species displaying very high polydispersity. Theoretical mass of rSslE is 160.0 kDa. The signal from the refractive index and light scattering are shown as a yellow and black line, respectively (left, y axis), and the average molecular weight calculated across each peak is shown as a red line (right, y axis).

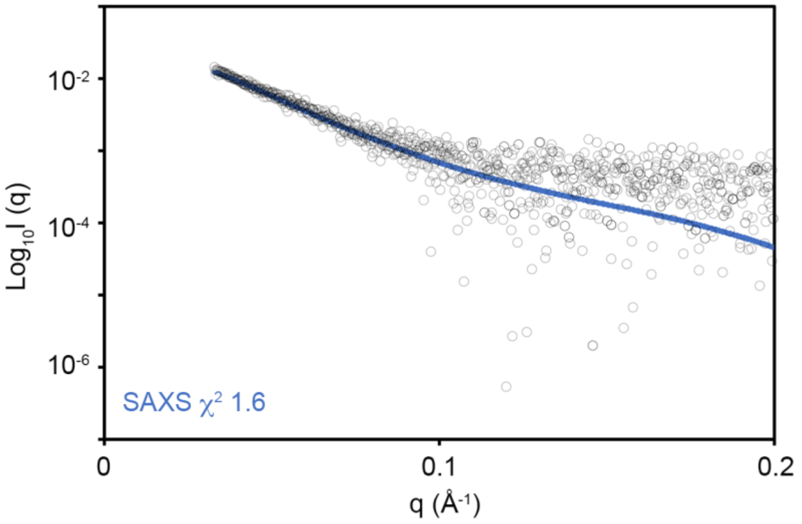


**Supplementary Fig. 17 | Fit of the rSslE fibre SAXS bead model to SAXS data.** Bead model is shown as blue line and SAXS data as black open circles, with $\chi^2$ of 1.6.

### Supplementary Tables

#### Supplementary Table 1: circular dichroism data processing.

| **Secondary structure type** | **% secondary structure** | | **Difference** |
|---|---|---|---|
| | pH 4.4 | pH 7.4 | |
| α-helix | 21.7 | 29.9 | 8.2 |
| β-sheet | 17.3 | 12.2 | 5.1 |
| β-turns | 14.2 | 13.1 | 1.1 |
| Random coil | 46.8 | 44.7 | 2.1 |
| NRMSD | 0.013 | 0.007 | - |

**Supplementary Table 2: SslE SAXS data and refinement statistics**

| | rSslE solution data | | COSMiCS fibrillation data | | rSslE fibre modelling |
|---|---|---|---|---|---|
| **SAXS data collection** | pH 7.4 | pH 4.4 | monomer | fibre | t=10 hrs |
| Beamline | DLS B21 | DLS B21 | DLS B21 | DLS B21 | DLS B21 |
| Wavelength (Å) | 1.0 | 1.0 | 1.0 | 1.0 | 1.0 |
| q Range ($Å^{-1}$) | 0.003 to 0.44 | 0.003 to 0.44 | 0.003 to 0.44 | 0.003 to 0.44 | 0.003 to 0.44 |
| **Structural parameters** | | | | | |
| I(0) | 0.203 ± 1.5e-04 | 0.206 ± 1.6e-04 | - | - | - |
| $R_g$ (nm) (from Guinier) | 4.03 ± 0.44 | 3.92 ± 0.13 | 3.92 | - | - |
| $R_g$ (nm) (from P(r)) | 4.07 ± 0.01 | 3.97 ± 0.01 | 3.98 | 51.1 | 4.34 ± 0.03 |
| $D_{max}$ (nm) (from P(r)) | 14.1 | 13.7 | 13.3 | 140 | 14.0 |
| Dammin model $\chi^2$ fit | 1.19 | 0.96 | - | - | 1.63 |
| Dammin models NSD (20 models) | 0.56 | 0.45 | - | - | 0.51 |
| EM model $\chi^2$ fit | 113.70 | - | - | - | - |
| **Molecular mass determination** | | | | | |
| MW (kDa) (from sequence) | 159986 | 159986 | 159986 | - | - |
| MW (kDa) (from SAXS) | 151106 | 158202 | 146800 (Bayesian Estimate) | - | - |

#### Supplementary Table 3: Primers, plasmids and strains used in this study

| Primer | Description | Sequence (5' to 3') |
|---|---|---|
| BD1 | Recombinant expression of *P. gingivalis* W50 RgpB-CTD F | GACGACGACAAGATGGGTACATCTATTGCCGACGTAG |
| BD2 | Recombinant expression of *P. gingivalis* W50 RpgB-CTD R | GAGGAGAAGCCCGGTTACTTCACTATAACCTTTTCTG TATACGTC |
| PC3 | Gene KO - *E. coli* JW5925-1 *sslE* downstream | TTATTTCATGCCGGATGCGGCGTGAACGCCTTATCCG GCATACAGGATTATGTAGGC |
| PC4 | Gene KO -*E. coli* JW5925-1 *sslE* upstream | TTTCTCCCAGTTACGAATTTTTAACATTTTGTCAGTT GCGTTATTAATTCCGGGGA |
| PC5 | *sslE* complementing plasmid F | CCGGGCTAGCTATCAATGATGTCGTTTTCTTAAGAAT GGA |
| PC6 | Full *sslE* complementing plasmid R | CCGGAAGCTTTTACTCGACAGACATCTTATGCTCGGT AACCT |
| PC7 | *sslEΔM60* complementing plasmid R | CCGGAAGCTTTTACGGGTTTCATCATCGCCGCTTTGCT G |

| Plasmid | Description | Reference |
|---|---|---|
| pPC1 | pOPINE expression of *E. coli* W SslE (full) residues 67-1497 | [8] |
| pPC2 | pET28b expression of *E. coli* SslE (NT1) residues 67-211 | This study |
| pPC3 | pET28b expression of *E. coli* SslE (NT2) residues 230-425 | This study |
| pPC4 | pET28b expression of *E. coli* SslE (NT1-NT2) residues 67-425 | This study |
| pPC5 | pET28b expression of *E. coli* SslE (NT3-M60) residues 426-1497 | This study |
| pBD1 | pET46 Ek/LIC expression of *P. gingivalis* W50 RpgB (CTD) residues 662-736 | This study |
| pOPINE | Expression vector | [9] |
| pET28b | Expression vector | Novagen |
| pET46 Ek/LIC | Expression vector | Novagen |
| pKD46 | Red recombinase expression | [10] |
| pBAD18-Cm | Complementation plasmid | [11] |
| pCPC1 | Full *sslE* complementing plasmid | This study |
| pCPC2 | *sslEΔM60* complementing plasmid | This study |

| *E. coli* strain | Serogroup/ genotype | Reference |
|---|---|---|
| DH5a | *F- Φ80lacZΔM15 Δ(lacZYA-argF) U169 recA1 endA1 hsdR17(rk-, mk+) phoA supE44 thi-1 gyrA96 relA1 λ-* | Invitrogen |
| BL21 (DE3) | *fhuA2 [lon] ompT gal (λ DE3) [dcm] ∆hsdS*<br>*λ DE3 = λ sBamHIo ∆EcoRI-B int::(lacI::PlacUV5::T7 gene1) i21 ∆nin5* | |
| Shuffle T7 | *fhuA2 lacZ::T7 gene1* [lon] *ompT ahpC gal λatt::*pNEB3-r1-*cDsbC* (Spec$^R$, *lacI*$^q$) ∆*trxB sulA11 R(mcr-73::miniTn10*--Tet$^S$)2 [dcm] *R(zgb-210::Tn10* --Tet$^S$) *endA1 Δgor* ∆*(mcrC-mrr)114::IS10* | NEB |
| JW5925-1 | F-, *Δ(araD-araB)567, ΔlacZ4787*(::rrnB-3), *λ*$^-$, *ΔyghJ747::kan*, *rph-1, Δ(rhaD-rhaB)568, hsdR514* | [12] |
| W (ATCC 35401) | Serotype (O6:K-:H49) CA$^+$ | [13] |

| | | |
|---|---|---|
| W Δ*sslE* | *sslE* gene disrupted with a $kan^R$ cassette | This study |
| H10407 (ATCC 35401) | Serotype O78:H11 | [14] |
| H10407 Δ*sslE* | *sslE* gene disrupted with a $kan^R$ cassette | This study |

#### Supplementary Table 4: Synthetic genes

| Gene | Description | Sequence (5' to 3') |
|---|---|---|
| gPC2 | *E. coli* SslE (NT1) residues 67-211 | CCATGGTCAAAACCGGTTATCTGACCTTAGGTGGTAGCCTGCGTGTTACCGGTGATATT<br>ACCTGTAATGATGAAAGCTCAGATGGCTTTACCTTTACACCGGGTGATAAAGTTACCTG<br>TGTTGCAGGTAATAATACCACCATTGCAACCTTTGATACCCAGAGCGAAGCAGCACGTA<br>GTCTGCGTGCAGTTGAAAAAGTTAGCTTTAGCCTGGAAGATGCACAAGAACTGGCAGGT<br>AGCGATAACAAAAAAAGCAATGCACTGAGCCTGGTTACCAGCATGAATAGCTGTCCGGC<br>AAATACCGAACAGGTTTGTCTGGAATTTAGCAGCGTTATTGAAAGCAAACGTTTCGATA<br>GCCTGTATAAGCAGATTGATCTGGCACCGGAAGAATTCAAAAAACTGGTTAATGAAGAA<br>GTGGAAAACAACGCAGCAACCGATAAAGCACTCGAG |
| gPC3 | *E. coli* SslE (NT2) residues 230-425 | CCATGGTCGATCTGAATGCAAGCTTTGTTAGCGCAAATGCCGAACAGTTTTATCAGTAT<br>CAGCCGACCGAAATTATTCTGAGCGAAGGTCGTCTGGTTGATAGCCAAGGTGATGGTGT<br>TGTTGGTGTTAACTATTACACCAATAGCGGTCGTGGTGTTACCGGTGAAAATGGTGAAT<br>TTAGCTTTAGCTGGGGTGAAACCATTAGCTTTGGTATTGATACCTTTGAACTGGGTAGC<br>GTGCGTGGTAATAAAAGCACCATTGCACTGACCGAACTGGGTGATGAAGTTCGTGGTGC<br>AAATATTGATCAACTGATCCACCGTTATAGCAAAGCAGGTCAGAATCATACCCGTGTTG<br>TTCCGGATGAAGTGCGTAAAGTTTTTGCAGAATATCCGAACGTGATCAACGAAATTATC<br>AATCTGAGCCTGTCAAATGGTGCAACCTTAGGTGAAGGTGAACAGGTTGTTAATCTGCC<br>GAACGAATTCATCGAACAGTTCAAAACCGGTCAGGCCAAAGAAATTGATACCGCAATTT<br>GTGCAAAAACCGATGGTTGTAATGAAGCACGTTGGTTTAGCCTGACCACACGTAATGTT<br>AATGATCTCGAG |
| gPC4 | *E. coli* SslE (NT1-NT2) residues 67-425 | CCATGGTCAAAACCGGTTATCTGACCTTAGGTGGTAGCCTGCGTGTTACCGGTGATATT<br>ACCTGTAATGATGAAAGCTCAGATGGCTTTACCTTTACACCGGGTGATAAAGTTACCTG<br>TGTTGCAGGTAATAATACCACCATTGCAACCTTTGATACCCAGAGCGAAGCAGCACGTA<br>GTCTGCGTGCAGTTGAAAAAGTTAGCTTTAGCCTGGAAGATGCACAAGAACTGGCAGGT<br>AGCGATAACAAAAAAAGCAATGCACTGAGCCTGGTTACCAGCATGAATAGCTGTCCGGC<br>AAATACCGAACAGGTTTGTCTGGAATTTAGCAGCGTTATTGAAAGCAAACGTTTCGATA<br>GCCTGTATAAGCAGATTGATCTGGCACCGGAAGAATTCAAAAAACTGGTTAATGAAGAA<br>GTGGAAAACAACGCAGCAACCGATAAAGCACCGAGCACACATACCAGTCCGGTTGTTCC<br>GGCAACCACACCGGGTACAAAACCGGATCTGAATGCAAGCTTTGTTAGCGCAAATGCGG<br>AACAGTTTTATCAGTATCAGCCGACCGAAATTATTCTGAGCGAAGGTCGTCTGGTTGAT<br>AGCCAAGGTGATGGTGTTGTTGGTGTTAACTATTACACCAATAGCGGTCGTGGTGTGAC<br>CGGTGAAAATGGTGAATTTAGCTTTTCATGGGGTGAAACCATTAGCTTTGGCATCGATA<br>CCTTTGAACTGGGTAGCGTTCGTGGTAATAAAAGTACCATTGCACTGACCGAACTGGGT<br>GATGAAGTTCGTGGTGCAAATATTGATCAACTGATCCACCGTTATAGCAAAGCAGGTCA<br>GAATCATACCCGTGTTGTGCCGGATGAAGTGCGTAAAGTTTTTGCAGAATATCCGAACG<br>TGATCAACGAAATTATCAATCTGAGCCTGAGCAATGGTGCAACCTTAGGCGAAGGTGAA<br>CAGGTTGTTAATCTGCCGAATGAATTCATCGAACAGTTTAAAACCGGTCAGGCCAAAGA<br>AATTGATACCGCAATTTGTGCAAAAACCGATGGTTGTAATGAAGCACGTTGGTTTAGCC<br>TGACCACACGTAATGTTAATGATCTCGAG |

| | | |
|---|---|---|
| gPC5 | *E. coli* SslE (NT3-M60) residues 426-1497 | CCATGGTCGGTCAGATTCAGGGTGTTATTAACAAACTGTGGGGTGTCGATACCAACTAT<br>AAAAGCGTTAGCAAATTTCACGTGTTCCACGATAGCACCAACTTTTATGGTAGCACCGG<br>TAATGCACGTGGTCAGGCAGTTGTGAATATTAGCAATGCAGCATTTCCGATTCTGATGG<br>CACGTAACGATAAAAACTATTGGCTGGCCTTTGGTGAAAAACGTGCATGGGATAAAAAT<br>GAGCTGGCCTATATTACCGAAGCACCGAGCATTGTTCGTCCGGAAAATGTTACCCGTGA<br>AACCGCAACCTTTAATCTGCCGTTTATTAGCCTGGGTCAAGTTGGTGATGGTAAACTGA<br>TGGTTATTGGTAACCCGCATTATAACAGCATTCTGCGTTGTCCGAATGGTTATAGCTGG<br>AATGGTGGTGTTAATAAAGATGGTCAGTGTACCCTGAATAGCGATCCGGATGATATGAA<br>AAACTTCATGGAAAATGTGCTGCGCTATCTGAGCAATGATCGTTGGCTGCCGGATGCAA<br>AAAGCAGCATGACCGTTGGCACCAATCTGGATACCGTTTATTTCAAAAAACATGGTCAG<br>GTGCTGGGTAATAGCGCACCGTTTGCATTTCATAAAGATTTTACCGGCATTACCGTTAA<br>ACCGATGACCAGCTATGGTAATCTGAACCCGGATGAAGTTCCGCTGCTGATTCTGAATG<br>GTTTTGAATATGTTACCCAGTGGGGTAGTGATCCGTATAGCATTCCGCTGCGTGCAGAT<br>ACCAGCAAACCGAAACTGACCCAGCAGGATGTTACCGATCTGATTGCATATATGAATAA<br>AGGTGGTAGCGTGCTGATTATGGAAAACGTTATGAGCAACCTGAAAGAAGAAAGCGCAA<br>GCGGTTTTGTTCGTCTGCTGGATGCAGCAGGTCTGAGCATGGCACTGAATAAAGTGTG<br>GTTAATAATGATCCGCAGGGTTATCCTGATCGTGTTCGTCAGCGTCGTAGCACCCCGAT<br>TTGGGTTTATGAACGTTATCCGGCAGTTGATGGCAAACCGCCTTATACCATTGATGATA<br>CCACCAAAGAAGTGATCTGGAAGTATCAGCAAGAAAACAAACCTGATGATAAGCCGAAA<br>CTGGAAGTTGCAAGCTGGCAAGAAGAAGTTGAAGGTAAACAGGTGACCCAGTTTGCCTT<br>TATTGATGAAGCAGATCATAAAACACCGGAAAGCCTGGCAGCAGCAAAACAGCGTATTC<br>TGGATGCATTTCCTGGTCTGGAAGTTTGTAAAGATAGCGATTATCACTATGAGGTGAAC<br>TGCCTGGAATATCGTCCTGGCACCGGTGTTCCGGTTACCGGTGGTATGTATGTTCCGCA<br>GTATACCCAGCTGGATCTGGGTGCCGATACCGCAAAAGCAATGCTGCAGGCAGCAGATC<br>TGGGCACCAATATTCAGCGTCTGTATCAGCATGAACTGTATTTTCGTACCAATGGTCGT<br>CAGGGTGAACGTCTGAATAGTGTGGATCTGGAACGCCTGTATCAGAATATGAGCGTTTG<br>GCTGTGGAACGAAACCAAATATCGTTATGAAGAGGGCAAAGAAGATGAGCTGGGCTTTA<br>AAACCTTTACCGAATTTCTGAACTGCTACACCAATAATGCATATGTTGGCACCCAGTGT<br>AGCGCAGAACTGAAAAAAAGTCTGATCGACAACAAAATGATCTACGGTGAGGAAAGCAG<br>CAAAGCCGGTATGATGAATCCGAGCTATCCGCTGAATTATATGGAAAAACCGCTGACAC<br>GTCTGATGCTGGGTCGTAGTTGGTGGGATCTGAACATTAAAGTTGACGTGGAAAAATAT<br>CCGGGTGCAGTTAGCGAAGAAGGTCAGAACGTTACCGAAACCATTAGCCTGTATAGCAA<br>TCCGACCAAATGGTTTGCAGGTAATATGCAGAGCACCGGTCTGTGGGCACCAGCACAGA<br>AAGAAGTTACCATTAAAAGTAATGCCAATGTGCCGGTGACCGTTACCGTTGCACTGGCA<br>GATGATCTGACCGGTCGTGAAAAACATGAAGTTGCACTGAATCGTCCGCCTCGTGTTAC<br>CAAAACCTATTCATTAGATGCAAGCGGCACCGTGAAATTCAAAGTTCCGTATGGTGGTC<br>TGATCTATATCAAAGGTGATAGCAAAGATAATGAGAGCGCCAGCTTTACCTTTACAGGT<br>GTTGTTAAAGCACCGTTCTATAAAGACGGTGCCTGGAAAAACGATCTGAATTCTCCGGC<br>ACCGCTGGGTGAACTGGAAAGTGATGCATTTGTTTATACCGCACCGAAAAAGAATCTGA<br>ACGCAAGCAATTATACCGGTGGCCTGAAACAGTTTGCAAATGATCTGGACACCTTTGCC<br>AGCTCCATGAATGATTTTTATGGTCGCAATGAAGAGGACGGCAAACATCGTATGTTTAC<br>CTATAAAAACCTGACGGGTCATAAACACCGTTTTGCCAATGATGTGCAGATTAGCATTG<br>GTGATGCACATAGCGGTTATCCGGTTATGAATAGCAGCTTTAGCACCAATAGCACCACA<br>CTGCCGACCACACCGCTGAACGATTGGCTGATTTGGCATGAAGTGGGTCATAATGCAGC<br>AGAAACTCCGCTGACCGTTCCGGGTGCCACCGAAGTTGCCAATAATGTTCTGGCACTGT<br>ATATGCAGGATCGTTATCTGGGTAAAATGAATCGTGTTGCCGATGATATTACCGTGGCA<br>CCGGAATATCTGGAAGAAAGCAATGGTCAGGCCTGGGCACGTGGTGGTGCGGGTGATCG<br>CCTGCTGATGTATGCACAGCTGAAAGAATGGGCAGAAAAAAACTTCGACATCAAAAAGT<br>GGTATCCGGAAGGCGAACTGCCGAAATTTTTCAGCGATCGTGAAGGTATGAAAGGCTGG<br>AACCTGTTTCAACTGATGCATCGTAAAGCCCGTGGTGATGATGTTGGTGATAAAACATT<br>TGGTGGCAAAAACTACTGTGCCGAAAGTAATGGTAATGCAGCTGATACCCTGATGCTGT<br>GTGCGAGCTGGGTTGCACAGACCGATCTGTCAGAATTCTTCAAAAAATGGAATCCTGGT<br>GCCAATGCATATCAGTTACCGGGTGCAAGCGAAATGAGCTTTGAAGGTGGTGTGAGCCA<br>GAGCGCATATAATACCCTGGCAAGCCTGAAACTGCCTAAACCTGAACAGGGTCCTGAAA<br>CAATCAACAAAGTTACCGAACATAAGATGAGCGTGGAACTCGAG |

*Restriction sites are underlined
